# Selective loss of diversity in doubled-haploid lines from European maize landraces

**DOI:** 10.1101/817791

**Authors:** Leo Zeitler, Jeffrey Ross-Ibarra, Markus G Stetter

## Abstract

Maize landraces are well adapted to their local environments and present valuable sources of genetic diversity for breeding and conservation. But the maintenance of open-pollinated landraces in *ex-situ* programs is challenging, as regeneration of seed can often lead to inbreeding depression and the loss of diversity due to genetic drift. Recent reports suggest that the production of doubled-haploid (DH) lines from landraces may serve as a convenient means to preserve genetic diversity in a homozygous form that is immediately useful for modern breeding. The production of doubled-haploid (DH) lines presents an extreme case of inbreeding which results in instantaneous homozygosity genome-wide. Here, we analyzed the effect of DH production on genetic diversity, using genome-wide SNP data from hundreds of individuals of five European landraces and their related DH lines. In contrast to previous findings, we observe a dramatic loss of diversity at both the haplotype level and that of individual SNPs. We identify thousands of SNPs that exhibit allele frequency differences larger than expected under models of neutral genetic drift and document losses of shared haplotypes. We find evidence consistent with selection at functional sites that are potentially involved in the diversity differences between landrace and DH populations. Although we were unable to uncover more details about the mode of selection, we conclude that landrace DH lines may be a valuable tool for the introduction of variation into maize breeding programs but come at the cost of decreased genetic diversity and increased genetic load.

## Introduction

Maize is an outcrossing species and has been cultivated for millenia in open-pollinated populations known as landraces. Mass selection in these populations has been highly successful, allowing maize landraces to adapt to a breadth of environments and a wide array of cultural preferences (Bellon *et al.* 2018). Over the last century, an inbred-hybrid system has replaced landraces in modern agriculture due to its higher yields and increased stability (Troyer 2001). But the inbred-hybrid system has focused on an ever-decreasing pool of germplasm, restricting genetic variation compared to landraces (van Heerwaarden *et al.* 2012).

Though lower-yielding in industrial conditions, landraces continue to serve as an important genetic resource for future crop improvement and adaptation (Sood *et al.* 2014; Gates *et al.* 2019). But the conservation of landrace diversity imposes a number of challenges. *In situ* conservation by practicing farmers has been very successful (Bellon *et al.* 2018), but is vulnerable to changing economic considerations and does not provide easy access for breeders. Conservation of germplasm *ex-situ* provides straightforward and safe long-term access to plant breeders, but genebank accessions have to be maintained as large populations to prevent the loss of diversity due to drift (Ellstrand and Elam 1993).

Recently, Melchinger *et al*. (2017) suggested using doubled-haploid (DH) technology as a means of conserving landrace diversity in a homozygous form that would simplify germplasm conservation and be more readily usable by plant breeders (Sood *et al*. 2014; Gorjanc *et al*. 2016; Melchinger *et al*. 2017; Mayer *et al*. 2017). The directed induction of DH lines has been developed for several crops to accelerate breeding (Smith *et al.* 2008; Gomez-Pando *et al.* 2009; Dunwell 2010). The technology permits the instantaneous development of homozygous lines within a single generation instead of six to ten generations of conventional recurrent self-pollination (Prigge *et al.* 2012). While not all landraces produce DH lines with equal success, Melchinger *et al*. (2017) concluded that genetically stable DH line libraries of landrace accessions could be used for *ex-situ* conservation of maize without major loss in genetic diversity.

Theoretical considerations, however, suggest that the instantaneous inbreeding associated with DH production may impact genetic diversity and fitness (Keller and Waller 2002; Charlesworth and Willis 2009). Inbreeding reduces the effective size of a population, resulting in increased genetic drift and a loss of genetic diversity (Charlesworth and Willis 2009). Inbreeding is also thought to impact deleterious alleles and overall genetic load. As an outcrossing species, maize harbors a substantial number of deleterious, partially recessive alleles (Yang *et al.* 2017), mostly maintained at low frequencies (Mezmouk and Ross-Ibarra 2014; Yang *et al.* 2017) likely at mutation-selection balance (Eyre-Walker and Keightley 2007). Depending on the population history, inbreeding can have opposing effects on deleterious alleles. On one hand, inbreeding exposes recessive deleterious alleles, which can then be purged from the population (Keller and Waller 2002; Henn *et al.* 2015). But purging of deleterious alleles is most efficient when inbreeding occurs gradually over several generations (Keller and Waller 2002), rather than instantaneously as expected during DH line production. Instead, instantaneous homozygosity likely decreases the efficacy of selection and increases genetic load. These processes likely contribute to the highly reduced efficiency of DH production in outcrossing maize landraces compared to modern breeding germplasm that has already experienced conventional inbreeding (Böhm *et al*. 2017; Melchinger *et al.* 2017).

Given these considerations, here we re-evaluate the effects of DH production in maize landraces. We quantify the changes in genetic diversity due to DH production and investigate the role of drift and selection in creating the observed patterns. Combining published genotype and phenotype data from a number of sources (Melchinger *et al*. 2017; Mayer *et al*. 2017; Brauner *et al*. 2018), we analyze and compare samples from five populations of European maize landrace accessions and their derived DH lines. In contrast to previous reports (Melchinger *et al*. 2017), we find that landrace genetic diversity is not fully captured by DH line libraries. Although we are unable to pinpoint the causes underlying allele frequency changes at individual outlier loci, we find evidence suggesting that selection against recessive deleterious alleles causes reduced genetic diversity in DH populations. We conclude that DH technology is not suited to conserve maize landraces and its use would result in the loss of potentially important diversity and increased genetic load in germplasm collections.

## Material and Methods

### Data Preparation

We used genomic data of five European maize landraces and their derived DH lines to study the effect of instantaneous homozygosity. For the landrace derived DH lines (DH) we used data from Melchinger *et al*. (2017) from a total of *n* = 266 individuals genotyped on the Illumina MaizeSNP50 BeadChip (Ganal *et al.* 2011). The genotypes were derived from five accessions: Bugard (BU, *n* = 36) from France, Gelber Badischer (GB, *n* = 59), Schindelmeiser (SC, *n* = 58) and Strenzfelder (SF, *n* = 69) from Germany, and Rheinthaler (RT, *n* = 44) from Switzerland. For the landrace population samples (LR) of the same accessions, we used data of *n* = 137 individuals (*n* = 22, *n* = 46, *n* = 23, *n* = 23, *n* = 23, respectively) from Mayer *et al*. (2017) that were also used in the analyses of Melchinger *et al*. (2017), genotyped on the 600k Affymetrix Axiom Maize Genotyping Array (Unterseer *et al.* 2014, Table S1). After combining the two datasets based on physical positions (AGPv2), we removed SNPs that were monomorphic across all accessions in the LR and DH. For all further analyses, we then used updated positions (AGPv4) for the SNPs obtained from supplementary data in Mayer *et al*. (2017). We also removed SNPs that violated Hardy-Weinberg equilibrium in all LR accessions using exact tests with mid-*p* adjusted *p >* 0.05, as well as low quality SNPs that matched the quality criteria ‘off-target variant’, ‘CallRateBelowThreshold’ and insertion-type SNPs of the Affymetrix Axiom 600k genotyping chip (classifications followed Table S6 in Unterseer *et al*. (2014); for details see Table S2 and supplementary note), removing a total of 83,011 SNPs. In total, 64,930 genotypes (on average 0.6%) in DH individuals remained heterozygous and were set as missing data. Finally, we removed sites that were missing in all individuals by filtering out sites with missing data above 0.99 using plink 1.9 (Chang *et al.* 2015). This resulted in 533,190 SNPs in the LR, 37,967 of which overlapped between the LR and DH. We refer to the smaller set of data hereafter as the ‘50k’ dataset.

Unless otherwise specified, all statistical analyses described below were conducted using R (R Core Team 2018).

### Phasing and Imputation

To project all 533,190 SNPs of the LR dataset onto the DH data, we used a two-step approach to phase and impute the data using BEAGLE 5.0 (Browning and Browning 2009) for each LR-DH combination separately. First, we phased and imputed the LR data, then we used this data as a reference to impute the DH lines. Parameters used for BEAGLE were `ne=100000 phase-states=200 nsteps=14`. To assess the quality of the imputation and establish optimal parameters for the algorithm, we dropped 10,000 known SNPs randomly in the DH and calculated imputation error rates for the *i*th SNP as 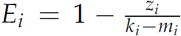, with *z* matches and *m* missing genotypes out of *k* individuals. The mean error rate was used to establish optimal imputation and phasing parameters for the algorithm after several runs with different parameters. We compared estimated error rates to diversity and recombination rate to exclude potential imputation biases. Estimated error rates varied from 10.6 % in GB to 15.9 % in BU, but are not correlated with recombination rate (Figure S1) and appear to be randomly distributed across the genome (Figure S2). Minor alleles are associated with higher error rates, however major alleles are mostly imputed correctly (Figure S3, see supplementary note). We refer to the set with 533,190 SNPs with imputed DH individuals and LR genotypes as the ‘600k’ dataset (Table S3).

### Genetic Analyses

To compare our findings to published results (Melchinger *et al.* 2017), we calculated nucleotide diversity (*π*) on a per-site basis for the 50k dataset using vcftools 0.1.17 (Danecek *et al.* 2011). We removed monomorphic sites within each LR-DH pair, calculated *π* of the remaining polymorphic sites within each landrace accession and compared *π* for these sites in the LR and DH using Mann-Whitney-Wilcoxon tests (Mann and Whitney 1947).

We used the R package SNPRelate (Zheng *et al.* 2012) to conduct a principal component analysis (PCA) for the 50k dataset to investigate the relationship between LR and DH. Furthermore, we calculated principal components in windows in a region around a putative inversion on chromosome 3 of the 600k LR set of accession BU using the R package lostruct (Li and Ralph 2019) with 500 SNPs per window and genome-wide using SNPRelate (Zheng *et al.* 2012).

We compared allele frequencies between DH and LR populations. First, we defined minor alleles using the pooled set of all DH and LR accessions and classified them as alternative alleles. Then, we calculated allele frequencies as counts of the alternative allele for each population using plink 1.9 (Chang *et al.* 2015).

We determined haplotypes and their respective frequencies in each population by concatenating SNPs in 50kb non-overlapping windows for the 600k dataset. After removing windows with ≤ 5 SNPs, haplotypes contained on average 21.2 SNPs. We repeated this analysis with windows based on genetic distance of 0.2 cM for the 50k data without filtering for short haplotypes due to low genomewide SNP density in the 50k data, resulting in windows with on average 8.7 SNPs. We identified the most abundant haplotype in each window in the LR, and classified haplotypes as ‘lost’, ‘fixed’ or ‘segregating’ according to their frequency in the DH. Haplotype diversity was calculated for all windows with at least two haplotypes in LR as 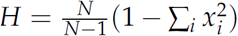) where *x*_i_ is the haplotype frequency determined in each bin and *N* the sample size, after Nei and Tajima (1981).

### Ancestral Allele Frequencies and Joint Probabilities

To distinguish between drift and selection as causes of allele frequency change during the process of DH production, we used a maximum likelihood grid search. We estimated ancestral frequencies and confidence intervals for the joint frequency spectra of DH and ancestral frequencies. Genotyped landrace individuals were sampled from the accession independently of those individuals that gave rise to the DH lines (Mayer *et al.* 2017; Melchinger *et al.* 2017). Because the landrace individuals are thus not the direct parents of the DH lines, a simple binomial sampling from the LR to DH to estimate confidence intervals would not be appropriate. Therefore, we considered for each accession three binomial sampling events in our estimation. From an ancestral landrace population, a first set of samples was genotyped (LR), and a second set used to produce DH lines from which another subsample was genotyped (DH) (Figure S4). For each accession and site of the 50k data, we estimated likelihoods across a grid of 100 possible ancestral frequencies ranging from to 0.99 as the product of these three binomial probability mass functions, defined as *P*(*k, n, p*), with *n* trials, *k* successes and *p* ∈ [0, 1] representing probabilities for three different sampling events, namely (1) the surviving DH lines from haploid induction until genotyping stage, created from the accession (*P*_*D*_), (2) the genotyped DH samples (*P*_*H*_) and (3) the genotyped landrace samples (*P*_*L*_).

For one accession this is a matrix with elements for the *j*-th ancestral frequency and *i*-th surviving DH individuals and can be estimated with

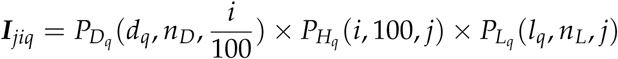

with *d* as the DH-allele count and *l* as the LR-allele count of the *q*-th site, and *n*_*D*_ and *n*_*L*_ as DH and LR chromosome counts. By maximizing the surface ***I***_*jiq*_ we obtain the maximum likelihood for the ancestral frequency’s probability *P*_*jiq*_.

Similarly, we computed a 95 % confidence interval by estimating a vector of probabilities for ancestral frequencies for the *s*-th DH line allele count and the *i*-th surviving DH individual by

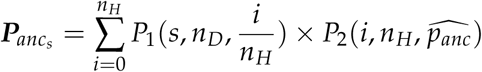

for each site (Figure S4). We used the central 95 % probability density of this distribution to define confidence intervals and defined SNPs outside of this confidence interval as allele frequency outliers (aSFS outliers).

We computed a second test statistic to infer outlier SNPs based on the joint probability for a given allele frequency in each population. Here, we computed the joint probability of landrace genotyping, DH line survival and DH line genotyping for each site. We model simple binomial sampling from an ancestral population with unknown allele frequency *x* which follows a beta distribution with parameters 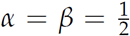. Integrating over this unknown frequency and using the notation above, the joint probability of observing *d* and *l* becomes:

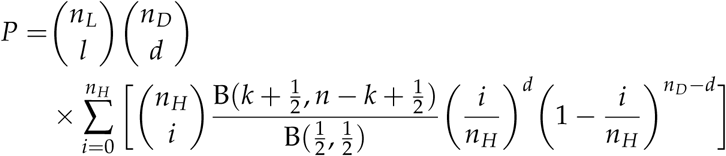

with *k* = *i* + *l* and *n* = *n*_*L*_ + *n*_*H*_ for each site and accession. We defined SNPs with joint probability in the top 5 % of the -*log*_10_(*P*) as outliers in each accession.

### Functional characterization of outlier SNPs

We hypothesized that outlier SNPs are deleterious and recessive and thus should be mostly found in heterozygous genotypes. To test this, we first asked whether outlier SNPs are deleterious and recessive by investigating their level of heterozygosity in the LR. We estimated genotype frequencies for all SNPs of the 50k dataset and removed non-outlier SNPs in linkage disequilibrium (*r*^2^ *>* 0.2) using plink 1.9 (Chang *et al*. 2015). We then compared equally-sized samples of outlier SNPs and non-outlier SNPs in 10 frequency bins between 0 and 1.

To study whether outliers affect functional phenotypic trait variation more than random sites, we computed trait effect sizes using a BayesB (Meuwissen *et al.* 2001) genomic prediction model implemented in GCTB (Zeng *et al.* 2018). We used arithmetic means over four locations of published phenotypes from 351 individuals of six European landrace DH line libraries (GB, RT, SF, Campan Galade, Walliser, Satu Mare) and 53 elite flint lines (Brauner *et al.* 2018). We calculated effect sizes based on the pooled dataset of these populations and additional parameters `–chain-length 30000 –burn-in 5000`. We used BEAGLE 5.0 (Browning and Browning 2009) to impute missing data after filtering based on the same cutoffs as the 50k dataset, resulting in 37,884 SNPs for 404 individuals. We then performed an ANOVA using absolute effect sizes for seven traits (shoot vigor, female flowering, *Fusarium* ear rot resistance, plant height, oil content, protein content and grain yield) separately with the following linear model:

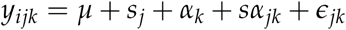

with *s* as the effect of the *j*-th SNP-type (outlier/non-outlier SNP), *α* as the effect of the binned (10 bins) *k*-th frequency in LR, *sα*_*jk*_ as the interaction effect between frequency bin and SNP-type, and *ϵ* as the residual effect.

To investigate the potential fitness consequences of outlier loci, we used published genomic evolutionary rate profiling (GERP) scores (Davydov *et al.* 2010; Rodgers-Melnick *et al.* 2015) estimated by Wang *et al*. (2017) from a multi-species whole-genome alignment of 12 plant genomes and the maize B73 reference genome (AGPv3) and corrected for reference genome bias (Wang *et al.* 2017). We used CrossMap 0.2.8 (Zhao *et al*. 2014) to update positions of GERP scores to version four of the maize B73 reference genome (AGPv4) (Jiao *et al.* 2017).

We compared outlier SNPs to an equally-sized set of non-outlier alleles stratified to match the ancestral allele frequency distribution of outlier SNPs by sampling from bins of 0.1 ancestral allele frequency. Then we calculated genetic distances in 1 Mbp windows based on a maize genetic map (Ogut *et al.* 2015) across the genome and partitioned SNPs into recombination quantiles. Lastly, we sampled equally sized fractions of outliers and non-outliers in recombination quantiles to accounted for differences in expected genetic load in genomic regions with different recombination rates. We calculated load based on observed genotypes under an additive and a recessive models. We then calculated the sum of overlapping GERP scores *>* 0 in 1 cM windows around each SNP for both models.

### Data and code availability

Scripts used for this project are stored in a public GitHub repository (https://github.com/LZeitler/eurodh-scripts).

## Results

### DH lines show decreased genetic diversity compared to landrace populations

We first evaluated the population structure of our samples using a genome-wide principal component analysis (PCA). Groupings largely follow overall expectations, with doubled-haploid (DH) and landrace (LR) individuals clustering well by accession on the first two principal components. The third principal component, however, separates a subgroup of the DH individuals from the main RT cluster (Figure 1A and S5).

**Figure 1.**
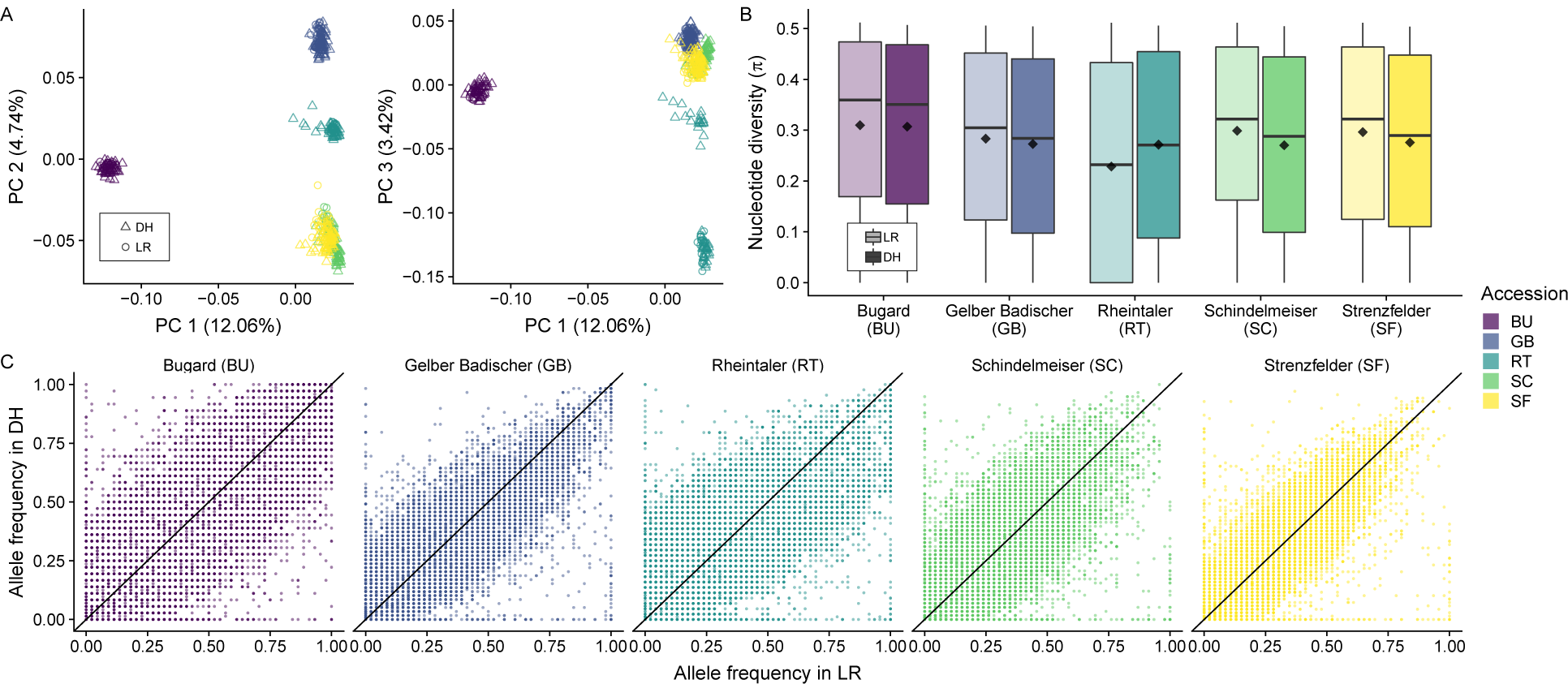
(A) Principal component analysis for the DH and LR of the 50k dataset. (B) Average per site nucleotide diversity at polymorphic sites for unimputed 50k data in each of the five LR-DH pairs. Means, represented by diamonds, are significantly different from each other within accessions (*p <* 1 × 10^−6^). (C) The joint site frequency spectrum (jSFS) for DH and LR populations. Allele counts are based on the published filtered dataset (Melchinger *et al.* 2017).

Using a set of genotyped SNPs (the 50k dataset, see Methods) from Melchinger *et al*. (2017), we compared per-site nucleotide diversity (*π*) in individual accessions (LR-DH pairs). We found that average *π* is significantly different between the LR and DH (two-sided Mann-Whitney-Wilcoxon *p <* 1 × 10^−6^ for all comparisons). In virtually all comparisons the nucleotide diversity is higher in LR than in DH (Figure 1B). In the accession RT, however, *π* is lower in the LR compared to DH, likely reflecting the population substructure observed among the DH (Figure 1A).

To investigate differences in diversity in more detail, we compared the joint site frequency spectrum (jSFS) of the two populations for each accession (Figure 1C). The jSFS shows substantial variation around the expected 1:1 line, highlighting differences in allele frequencies between the two populations. In particular, the jSFS reveals a number of alleles segregating in the LR that have been lost in the DH lines, as well as a smaller number that have been fixed. In some cases, we found segregating SNPs in the DH lines that were monomorphic in the respective LR population, suggesting that the frequency of the minor allele in the ancestral population must have been low and was not sampled in the genotyped set of LR individuals. Comparison of our jSFS with results from Melchinger *et al*. (2017) reveals striking differences (Figure S6B), as these authors appear to have filtered out a substantial proportion of alleles at low frequency in one or both populations, removing much of the observed signal of allele frequency change.

### Genome-wide pattern of allele frequency distortion

The jSFS highlights the dramatic difference in allele frequency between LR and DH populations. But, because DH lines were not the direct offspring of genotyped LR individuals (Figure S4), a direct comparison between LR and DH populations is not straightforward. To circumvent this difficulty, we first inferred allele frequencies in the ancestral population of both LR and DH (see Methods). Under neutrality, the expected allele frequency in the DH population is identical to that of the ancestor, and comparison of the site frequency between these populations (ancestral site frequency spectrum test; aSFS test) allows identification of alleles with unusual shifts in frequency (Figure 2). The number of outlier SNPs in the aSFS test (those outside of the 95% confidence interval around the ancestral frequency) varied between 1,769 (4.66%) in accession SF and 6,364 (16.78%) in accession RT. In total, we identified 12,345 distinct outlier loci in the five accessions. Of these, 9,305 were detected in only one of the accessions and 1,877, 539, 317, 307 SNPs overlapped in 2, 3, 4, 5 accessions, respectively (Table S4). A substantial fraction (14.71 %) of these outlier loci were segregating in LR but lost in DH, while only a very small minority of outlier loci (1.45 %) were fixed during DH production. In addition, sites that were outliers in multiple accessions had a higher mean frequency in LR and lower mean frequency in DH populations compared to outliers that were unique to one population (Figure S7A-B). Similarly, we found that a frequency reduction was more common in aSFS outliers shared among accessions while on average unique outliers changed only little (Figure S7C).

**Figure 2.**
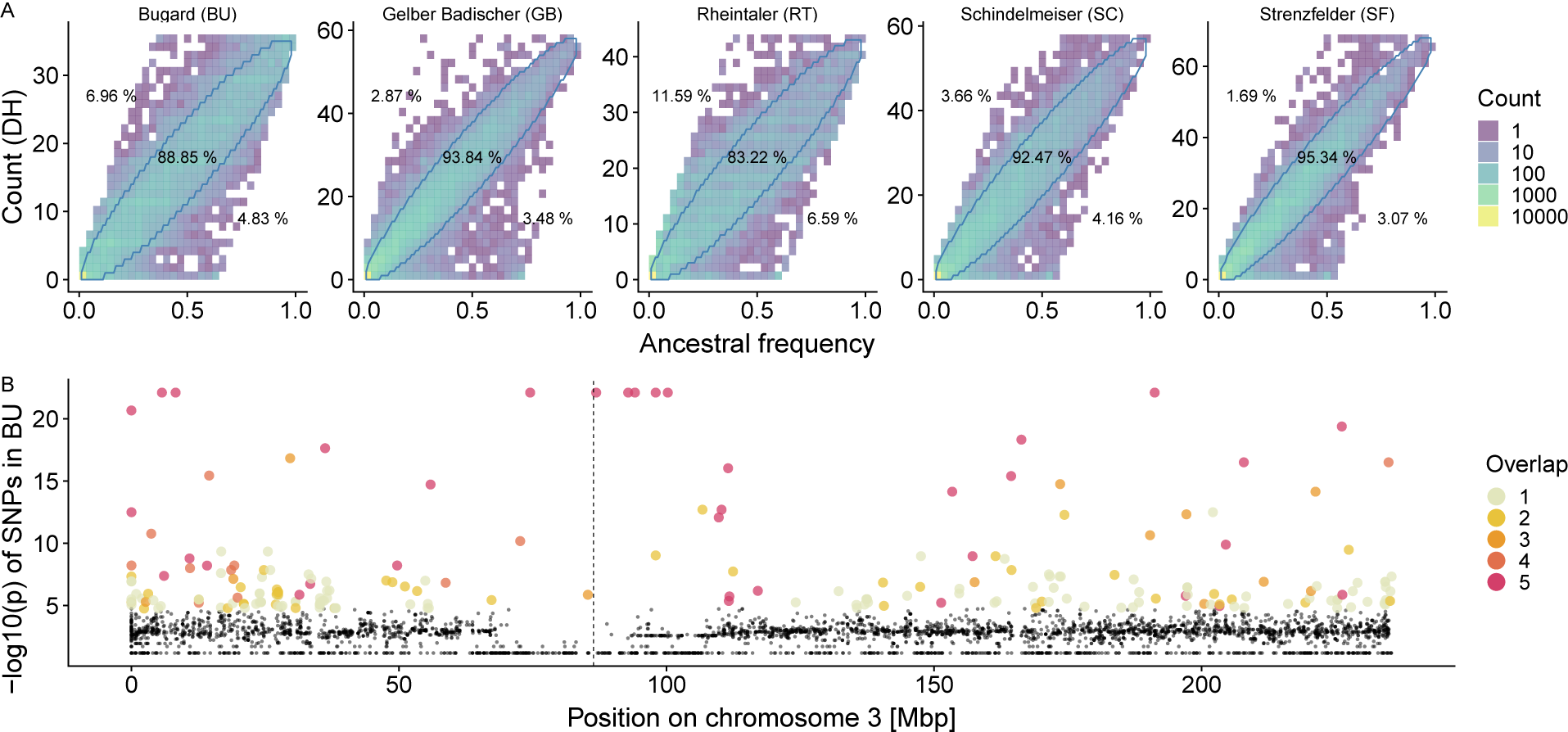
(A) Estimated ancestral and DH allele frequencies for all accessions of the 50k dataset show significant outliers, with the 95 % confidence interval represented by blue lines in the joint frequency spectrum (aSFS test). Percentages indicate the proportion of SNPs above, below and inside the interval. (B) Joint probability test along chromosome 3 in the Bugard landrace (BU). Colored dots represent the top 5 % – *log*_10_(*p*)-values which we defined as outliers. Colors represent the number of accessions in which a given locus is an outlier. The dashed line indicates the centromere position.

As a second means to identify outlier loci, we calculated the joint probability of the LR and DH genotypes (see Methods), yielding a *p*-value for each SNP (Figure S8). The test identified 9,458 outliers across accessions. We compared these outliers among different accessions and found that highly significant outlier SNPs were often shared among several populations (Figure S9).

The two approaches yielded largely similar results. Although the aSFS test identified nearly twice as many outliers as the joint probability test, virtually all of the outliers (97 %) identified in the joint probability test were also found in the aSFS comparison (Figure S10). Some clusters of co-located outlier sites were evident in both tests (Figure 2B). For instance, we detect a large region close to the centromere of chromosome three with multiple loci that were detected as outliers in multiple accessions.

### Haplotype tests reveal selection hotspots

Tests based on individual SNPs yielded outliers and indicated that these are often shared between accessions. However, such tests are limited in their power to detect changes in low frequency alleles. Haplotype based comparisons are more sensitive to rare alleles, as every new allele creates an additional haplotype. We imputed the 600k SNPs in the DH data using genotypes from the LR populations, and identified haplotypes in non-overlapping 50kb windows.

We observed a reduction in the number of haplotypes in the DH compared to the LR population: in the 600k data, we identified on average 7.65 segregating haplotypes per window in the LR and 4.36 in the DH. Haplotype diversity was significantly reduced in the DH compared to the LR populations (0.40 compared to 0.63; Figure 3A). The difference in haplotype diversity between LR-DH pairs was less pronounced in the 50k dataset, but haplotype diversity of the DH populations was also reduced in every accession compared to the LR (Figure S11 and S12A).

**Figure 3.**
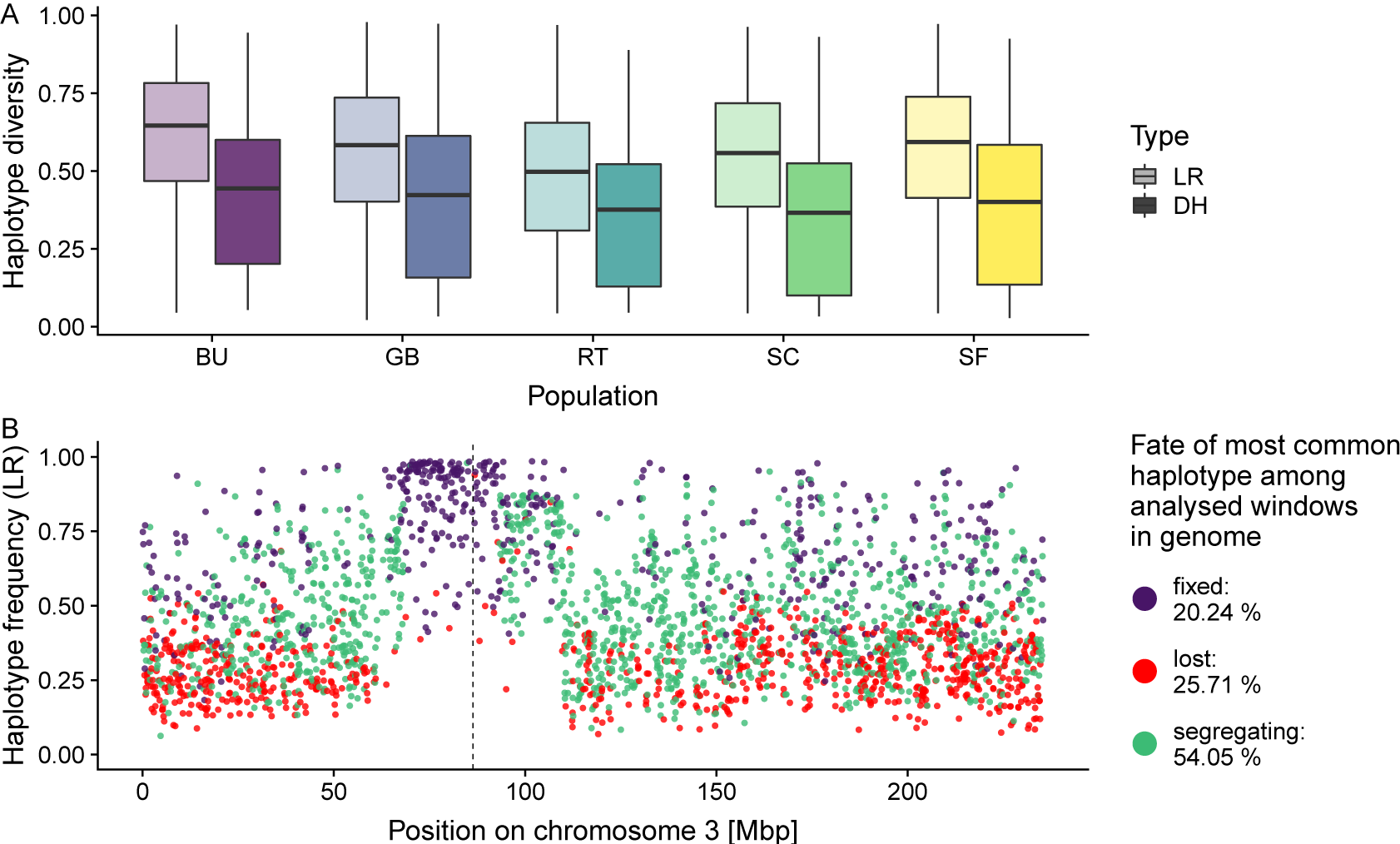
(A) Comparison of average haplotype diversity in 50kb windows for the 600k dataset in different accessions. Haplotype diversity of the imputed DH dataset is reduced compared to the LR dataset. (B) LR haplotype frequency in BU along chromosome 3, colored by the fate of the haplotype in the DH population. The centromere is shown as a vertical dashed line. Percentages listed in the legend correspond to genome-wide proportions of fixed, lost and segregating haplotypes.

We tracked the frequency change of the most common haplotype in each window between LR and DH populations in the 600k dataset and classified haplotypes according to their fate in the DH. While the majority (13,607 or 58.35 %) of the most common haplotypes in the LR were still segregating in the DH populations across accessions, a substantial minority (5,113, 21.93 %) were lost and another large fraction (4,600, 19.72 %) were fixed (Figure S13, Table S5).

Similar to the SNP outlier tests, we found several genomic regions with multiple consecutive windows exhibiting fixation or large changes in haplotype frequency (Figure S13). Windows with losses of major haplotypes coincided with highly significant joint probability outliers (Figure S14). In particular, the same region of chromosome 3 identified by SNP outliers showed strong signals of haplotype change in BU and RT, where even haplotypes with intermediate frequencies in LR were fixed in the DH (Figure 3B). To further investigate this region, we conducted a local PCA (Li and Ralph 2019) in the BU landrace data between positions 65 Mb and 95 Mb, revealing three distinct clusters across multiple consecutive windows (Figure S15A). This contrasts with genome-wide PCA (Figure S15B), and is consistent with previous reports of a segregating inversion polymorphism in this region (Romero Navarro *et al.* 2017).

To exclude potential effects of imputation errors, we performed the same analysis with the 50k data and windows based on genetic distance. The estimated fraction of fixed alleles was reduced and the putative inversion on chromosome 3 was no longer detected (Figure S12B), likely due to lower SNP density and imputational quality for minor alleles (Figure S3, supplementary note). Nonetheless, lost alleles occurred at similar rates (10.03 %, Figure S12B), confirming the results of the 600k data.

### Outliers are more heterozygous than random alleles

To characterize potential changes due to selection, we first asked whether outlier SNPs are more likely to be recessive compared to random sites. We hypothesized that recessive deleterious sites should show higher observed heterozygosity in the outbred LR as selection should effectively remove homozygous genotypes. Indeed, outliers had a significantly higher frequency of heterozygotes in the LR compared to a frequency-matched sample of non-outliers for all populations (*p <* 0.05; Figure 4 and S16).

**Figure 4.**
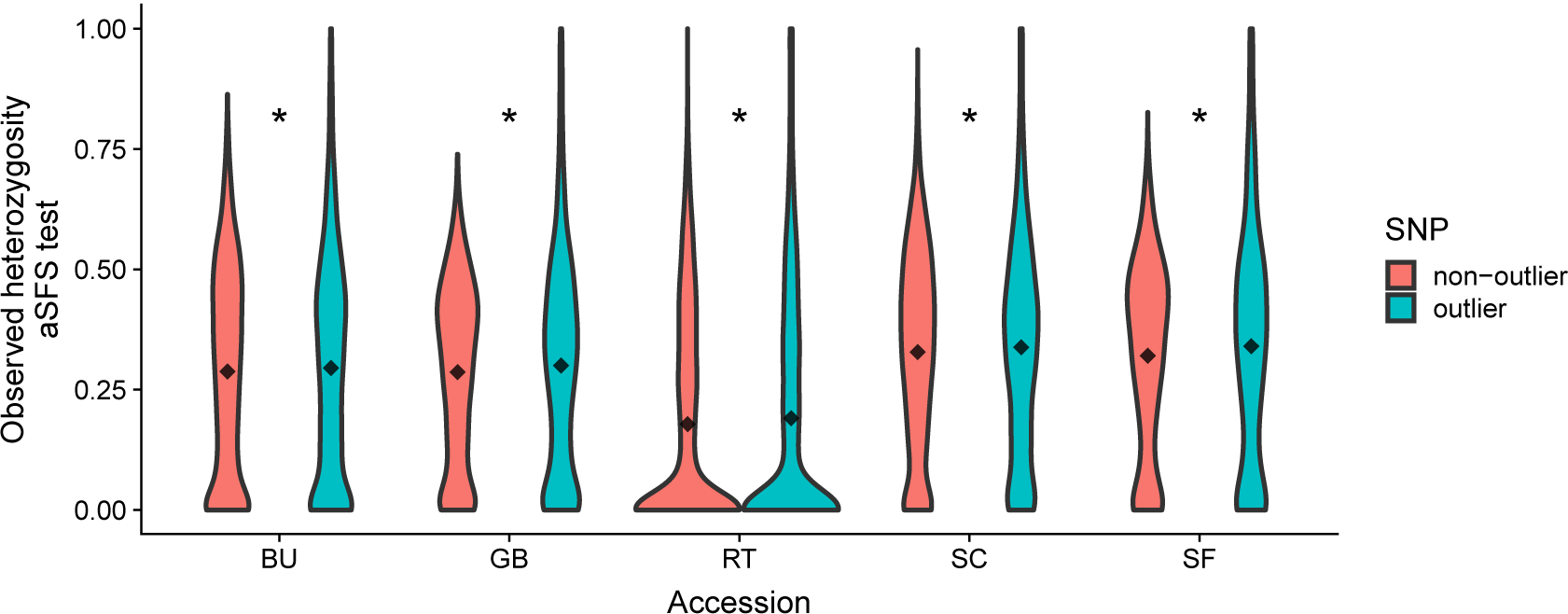
Violin plots for the frequencies of heterozygous genotypes of LD-pruned non-outlier SNPs and outlier SNPs in the LR accessions for aSFS outliers. Comparisons with asterisks have significantly different means (1,000 bootstraps, *p <* 0.05).

Next, we investigated whether outlier regions were enriched for functional variants. To test this, we estimated polygenic effect sizes for seven traits from a DH line panel with 404 individuals from different landraces using a BayesB prediction model (Meuwissen *et al.* 2001). We then tested whether effect sizes differed between outlier and non-outlier windows across a range of allele frequencies. Allele frequency was highly significant for all traits, and while outliers had significantly different absolute effect sizes for only two traits (plant height, oil content) we found significant interactions between SNP-type and allele frequency for all traits (Table S6). The relationship between effect size and allele frequency was negative for outlier SNPs, but positive for non-outlier SNPs across all traits after fitting linear regression models (Figure S17).

Lastly, we aimed to characterize changes in genetic load due to selection using published GERP estimates of evolutionary constraint at each SNP calculated from a phylogeny of 13 species (Wang *et al.* 2017). Previous studies in maize have shown that GERP scores correlate with estimated SNP effects on yield and are thus a quantitative proxy for the fitness effects of a locus (Yang *et al.* 2017). We analyzed whether outliers contribute higher genetic load than random SNPs by summing GERP scores in the 1 cM centered around each SNP. Under a recessive model, in four out of the five LR and DH populations outlier windows showed lower genetic load than random windows, while there was no significant difference in BU in both the LR and DH (t-test, *p* = 0.05; Figure S18). Under an additive model, the mean load of outliers was significantly lower compared to non-outlier in all accessions in the LR and in all but BU in the DH (t-tests, *p* = 0.05; Figure S18).

## Discussion

### Reduction in genetic diversity between landrace and DH populations

We find a significant reduction in genetic diversity during DH production in four out of five accessions (Figure 1B and 3A). The increase in *π* seen in RT may have resulted from the observed population sub-structure in the DH population, perhaps due to the use of seed from distinct rounds of regeneration *ex situ* (Chebotar *et al.* 2003). The RT landrace itself also exhibited higher homozygosity than samples of other LR populations, suggesting a history of inbreeding during conservation. Whatever the cause, our estimates of diversity at the haplotype level reinforce these overall findings, showing even greater losses of diversity than seen at individual SNPs (Figure 3A and S11). Altogether, these results closely follow theoretical predictions regarding the consequences of inbreeding (Charlesworth and Willis 2009; Schnable and Springer 2013).

The loss of diversity we observe in DH populations stands in contrast to previous findings (Melchinger *et al.* 2017) using the same data. For a detailed comparison, we reconstructed the jSFS using the original data from the previous study, revealing that Melchinger *et al*. (2017) had filtered the data in such a way as to remove sites with extreme allele frequencies in either population (Figure S6B, Melchinger *et al.* 2017). While minor allele frequency filters are often applied in quantitative genetic studies, the removal of rare alleles can strongly influence results of population genetic analyses (Weale 2010; Linck and Battey 2019). Moreover, such alleles are of particular interest for the conservation of genetic diversity. Therefore, we limited our filtering to data quality but did not remove rare alleles (Table S2).

### DH production creates selection hotspots

To understand the effect of DH line production on the change in diversity across the genome, we employed two outlier tests comparing the allele frequency changes between the LR and DH. These tests identified loci for which the allele frequency shifted more than expected by random drift. Outliers exist in all five European landrace accessions (Figure 2A). While the aSFS test resulted in a larger set of outliers than the joint probability test, both tests identified a largely overlapping set of outliers (Figure S10). And while many outliers were shared among accessions (Figure 2B and S9), indicating some shared signal resulting from DH production, the majority of outliers were accession specific. In other crops like potato, it has been shown that the genomic signals of inbreeding are largely specific to individual lines (Zhang *et al.* 2019). The increased strength of selection due to the instantaneous homozygosity during DH production and the shared history of European maize landraces might have caused the increased signal of shared outliers among accessions.

While we found outliers distributed across the whole genome, we also observed clustering in specific genomic regions (Figure 2B). The distribution of the fate of major haplotypes in windows along the genome revealed regions enriched for outliers that go to fixation or loss (Figure 3 and S11). One of the most pronounced signals was on chromosome 3 in BU and in RT. This approximately 25-Mb region (70-95 Mb) overlaps with a previously identified putative 6-Mb inversion that is associated with flowering time in maize (Romero Navarro *et al.* 2017). Further testing using local principle components indicated the presence of this inversion in the landrace sample of accession BU (Figure S15). In this landrace from Southern France, the inversion may be involved in flowering time adaptation. Alternatively, unconscious selection on flowering time might have occurred during haploid induction to synchronize landrace flowering with the inducer line or subsequent cultivation in northern latitudes. Other regions where outliers clustered in longer windows in all accessions (Figure S13) were mostly located in low recombination regions around centromeres (Ogut *et al.* 2015). Weakly deleterious alleles are likely to accumulate in such regions (Rodgers-Melnick *et al.* 2015; Yang *et al.* 2017), and if most fitness-affecting mutations are at least partially recessive (Yang *et al.* 2017) such regions might be expected to show selection when made homozygous during DH production.

Imputation of the 600k data strongly increased the marker density and therefore the power of our study. Yet, minor alleles were more affected by imputational error (Figure S3), which might introduce a potential bias towards a lower number of estimated haplotypes per window and therefore on haplotype diversity. Hence, we also performed the haplotype analysis using the 50k dataset. Despite the lower power due to the sparse marker density, the 50k data showed mostly similar patterns for haplotype diversity and major haplotype change. One difference was the lack of evidence for the putative inversion (Figure S12B). While the 600k data was designed with a more diverse sample of maize lines that included both haplotypes at this inversion, the region on the MaizeSNP50 chip does not include SNPs segregating for the inverted haplotype (see Pyhäjärvi *et al.* 2013). Low frequency polymorphisms should have only little effect on the fate of the most abundant haplotype during DH production, except for fixing major haplotypes in windows where all haplotypes were rare in the LR and minor alleles were incorrectly imputed. The higher –*log*_10_(*p*) values, i.e., outliers (based on unimputed data) overlapping with lost major haplotypes confirm that the fate of the major haplotype was little affected by imputation errors (Figure S14). We expect that most rare alleles in the DH should have been correctly imputed as we used the LR that they were derived from as reference for the imputation algorithm. In the absence of selection, alleles in the DH populations should exist at similar frequencies in their respective LR. Hence, potential imputation errors rather lead to an underestimation of the selective loss of alleles.

### Differences between DH and LR are potentially due to recessive deleterious load

Doubled-haploid lines show particularly poor fitness compared to outcrossing lines and even compared to inbred lines (Strigens *et al.* 2013; Böhm *et al.* 2014). The observed inbreeding depression in DH and inbred lines in maize is likely due to accumulation of deleterious alleles as a result of inbreeding (Bataillon and Kirkpatrick 2000; Charlesworth and Willis 2009). Recent work has shown that the observed decrease of heterozygosity during inbreeding of maize landraces is slower than expected, suggesting that the exposure of recessive deleterious alleles removes certain haplotypes and maintains heterozygosity (Roessler *et al*. 2019). While multiple cycles of inbreeding allow for recombination and purging of genetic load, DH production induces instantaneous homozygosity which reduces the possibility for effective purging. Moreover, heterozygosity can be regenerated by intercrossing DH lines only in regions of the genome where multiple haplotypes were captured during DH production. Our results show that this is not the case in >20% of the genome, where even the previously most common haplotype was lost (Figure 3B and S12B). For lower frequency haplotypes this effect is expected to be even more severe. Hence, a substantial number of haplotypes will not be available for further selection in intercrossed DH line populations.

Consistent with the assumption of recessive deleterious mutations, we find evidence that outlier SNPs are more likely to be heterozygous in LR populations (Figure 4 and S16). While outlier sites show strong shifts in allele frequency, they are unlikely to be directly causal themselves, but instead linked to deleterious sites. The comparison of genetic load around outliers and non-outliers, identified significantly lower load around outlier SNPs (Figure S18), which might indicate that the causal deleterious SNP in the region was not genotyped in our analysis. The relatively low marker density of the SNP chips, in addition to the ascertainment bias towards inbred breeding material, likely prevented assaying the causal deleterious alleles, but allowed us to capture the loss in diversity around causal sites.

Finally, we searched for evidence that outlier loci were particularly likely to contribute to phenotypic variation. We find evidence of selection in plant height and oil content (Table S6), despite our relatively simple additive GWAS model and the fact that most loci show at least partially recessive effects on yield (Yang *et al*. 2017). Negative slopes (Figure S17) in the regression models of allele frequency and absolute effect size are consistent with the deleteriousness of outlier SNPs and match previous findings in maize that deleterious mutations explain a significant proportion of phenotypic variance (Yang *et al.* 2017). The significant effect of outliers on plant height effect sizes might be the signature of natural or unintended selection on height or a related phenotype in maize landraces. While oil content is sometimes used as a means to identify haploid seed in the creation of DH lines (Prigge and Melchinger 2012), a different approach was used to create the DH lines used here (Melchinger *et al.* 2017), and we have no explanation for the signal of selection on oil content observed here.

Overall, our results suggest that the observed reduction in diversity within different populations is not caused by a few large-effect loci, but rather by a polygenic effect of partially recessive, mildly deleterious mutations (Bataillon and Kirkpatrick 2000).

### Conservation of landrace diversity

Landraces are an invaluable source of adaptive diversity (Bellon *et al.* 2018; Gates *et al.* 2019), and their conservation should remain a high priority for future generations. Here, we showed that DH line libraries from landraces do not capture the full diversity present in the landrace. Therefore, while DH line libraries present a valuable tool to introgress known alleles into breeding programs, we conclude they can not replace *ex situ* and *in situ* conservation efforts. To preserve landraces and their full genetic diversity, they should be reproduced in large populations to prevent inbreeding and the consequent shift of allele frequencies. An improved understanding of inbreeding and the underlying genomic changes it induces will help to conserve these genetic resources and harness their diversity to breed improved crop varieties.

## Acknowledgments

We thank Graham Coop, Michelle Stitzer and members of the Ross-Ibarra lab for helpful ideas and suggestions and Tobias Schrag and Albrecht Melchinger for supplying unfiltered DH line data. LZ was supported by SKH Carl Herzog von Württemberg, KWS Saat SE and The Ministry of Science, Research and the Arts of the State of Baden-Württemberg (Baden-Württembergisches Ministerium für Wissenschaft, Forschung und Kunst). JR-I was supported by NSF grant 1546719 and USDA Hatch project CADPLS2066H. MGS was supported by the Deutsche Forschungsgemeinschaft (DFG) grant STE 2654/1-1 and under Germany’s Excellence Strategy EXC-2048/1 Project ID 390686111. We would also like to thank Felix Andrews for statistical advice, although we did not follow it.

## Supplements

### Data preparation and quality control

Both datasets were combined based on reference genome V2 positions into the hapmap format. For SNPs where the opposite strand was targeted by the two array platforms the corresponding alleles were converted to their complementary basepair and compared to the landrace population reference. We removed insertions, unmapable SNPs (chr0 and duplicated SNPs), non-polymorphic sites, and SNPs with quality classes ‘off-target variant’ and ‘call rate below threshold’. We further removed SNPs that violated the Hardy-Weinberg equilibrium (*χ*^2^, *mid – p <* 0.05) in at least one LR accession from all LRs using plink 1.9 (Chang *et al*. (2015), see Table S2). A vcf file for the whole dataset was constructed using TASSEL 5 (Bradbury *et al.* 2007). Accession filtered datasets were written using custom R scripts with various packages.

### Imputation allows haplotype analysis

In our study we relied on published genotyping data based on two different genotyping arrays (Ganal *et al*. 2011; Unterseer *et al.* 2014). While the data was highly consistent and reconstructed the population structure correctly (Figure 1A and S5), these platforms come with several limitations, reducing the ability to detect rare and potentially deleterious alleles and reductions of diversity. By imputing the DH dataset we combined phase information of both the LR populations and the DH lines and were able increased the SNP density for the DH data. The imputed dataset enabled us to identify major reductions of mean haplotype diversity in polymorphic windows and large regions of complete fixation in the DH populations. The extent of this loss of diversity could only be detected using imputed genotypes. While imputation came with an error rate of 10.6 % to 15.9 %, we were able to increase the number of sites in the DH lines from 37 thousand to over 530 thousand. Imputational error rates depend highly on the reference panel used, MAF, SNP density and chromosome sample size; error rates in the literature range from 1 % to 15 % (Browning and Browning 2009; Howie *et al*. 2009; Khatkar *et al.* 2012). We show that the estimated imputational error rate is randomly distributed across the genome (Figure S2) and the correlation with the genetic distance of neighboring SNPs is low (Figure S1). Furthermore, the mean haplotype diversity of the unimputed 50k dataset (Figure S11) showed significant reductions of DH diversity compared to the LR in every accession and we found corresponding outliers using imputed and unimputed data (e.g., chromosome 3, near centromere). Additionally, we reanalyzed the fate haplotypes of the unimputed 50k and used windows based on LD (window size 0.2 cM). This resulted a reduction of fixed major haplotypes, but in similar amounts of lost major haplotypes confirm our findings of the 600k data run. Hence, we conclude that the trade-off between maker density and imputation error is justified for the haplotype analysis, and the information gain associated with imputation overcomes the loss in statistical power due to undetected genetic diversity in the DH. While genotyping arrays have high genotyping accuracy for called SNPs, future studies should use genome-wide sequencing to avoid imputation and ascertainment issues. This would allow harnessing the full potential of DH lines from landraces to study the causes of inbreeding depression in maize.

**Table S1.** Sample sizes for DH and LR

| Population | DH | LR | Sum |
| --- | --- | --- | --- |
| BU | 36 | 22 | 58 |
| GB | 59 | 46 | 105 |
| RT | 44 | 23 | 67 |
| SC | 58 | 23 | 81 |
| SF | 69 | 23 | 92 |
| Sum | 266 | 137 | 403 |

**Table S2.** Number of SNPs removed during quality control

| stage | removed |
| --- | --- |
| duplicated SNPs | 389 |
| Chromosome 0 | 310 |
| non polymorphic sites | 77798 |
| Insertions | 107 |
| quality tag: CallRateThresh | 846 |
| quality tag: off-target variant (OTV) | 2747 |
| violated HW | 814 |
| **SUM removed SNPs** | **83011** |

**Table S3.** Datasets used in this study

| Dataset | Populations | SNPs | Individuals |
| --- | --- | --- | --- |
| 50k DH | BU, GB, RT, SC, SF | 37,967 | 266 |
| 50k LR | BU, GB, RT, SC, SF | 37,967 | 137 |
| 600k DH | BU, GB, RT, SC, SF | 533,190 | 266 |
| 600k LR | BU, GB, RT, SC, SF | 533,190 | 137 |
| GWAS DH | CG, EF, GB, RT, SF, SM, WA | 37,884 | 404 |

**Table S4.**
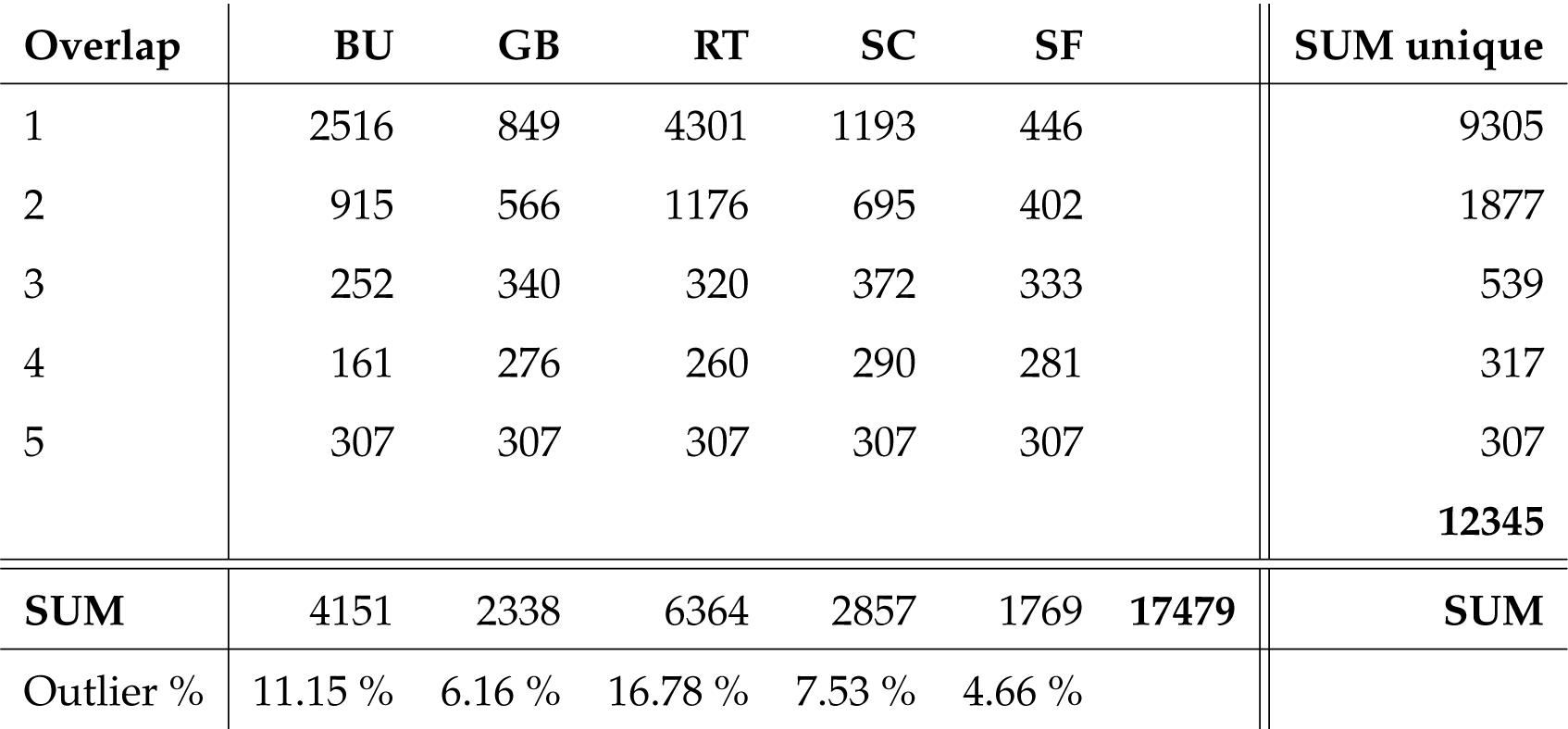
Number of outlier SNPs identified in the aSFS test

**Table S5.** Fate of most common haplotypes in a total of 34,833 50kb windows in the 600k data.

| Accession | fixed | lost | segregating | Sum (windows) | Sum (haplotypes) | fixed % | lost % | segregating % |
| --- | --- | --- | --- | --- | --- | --- | --- | --- |
| BU | 3774 | 6723 | 13186 | 23683 | 209982 | 15.94 | 28.39 | 55.68 |
| GB | 3222 | 3917 | 16684 | 23823 | 244164 | 13.52 | 16.44 | 70.03 |
| RT | 5999 | 3682 | 12339 | 22020 | 113418 | 27.24 | 16.72 | 56.04 |
| SC | 5199 | 5138 | 12993 | 23330 | 160030 | 22.28 | 22.02 | 55.69 |
| SF | 4806 | 6106 | 12837 | 23749 | 194009 | 20.24 | 25.71 | 54.05 |

### Additional figures

**Figure S1.**
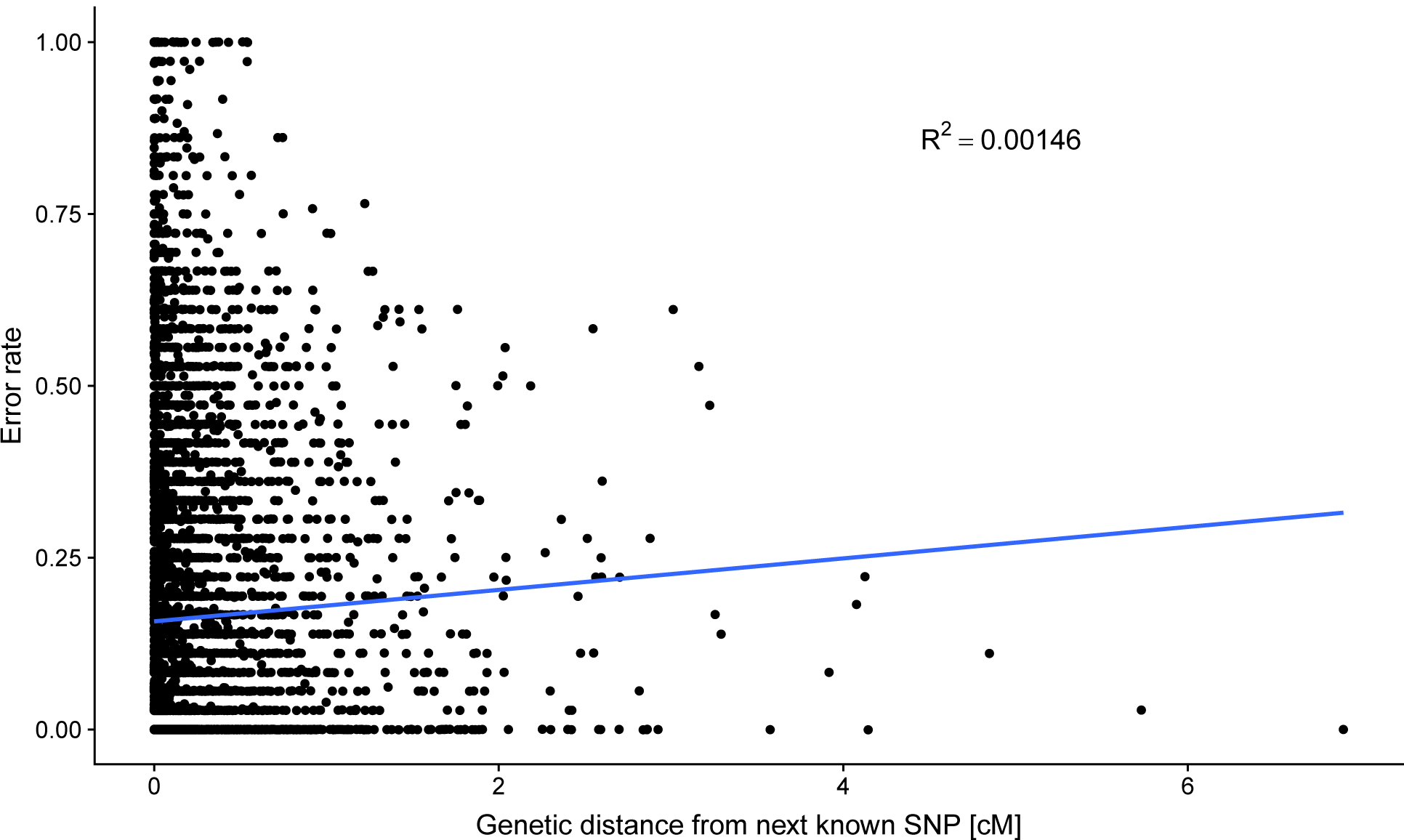
Correlation of marker density and imputation error rate shows low *R*^2^ (*R*^2^ = 0.00322).

**Table S6.** ANOVA tables of the outlier characterization using GWAS effect sizes. The term ‘outlier’ refers to the SNPs classified as ‘outlier’ or ‘non-outlier’ in the aSFS.

| Trait | Term | df | SumSq | MeanSq | F-value | p-value |
| --- | --- | --- | --- | --- | --- | --- |
| shoot vigor | outlier | 1 | 1.65e-07 | 1.65e-07 | 1.57 | 0.21 |
|  | frequency bin | 9 | 4.89e-05 | 5.44e-06 | 51.8 | 1.45e-94 |
|  | outlier:frequency bin | 9 | 1.34e-05 | 1.48e-06 | 14.2 | 4.26e-23 |
|  | Residuals | 113000 | 0.0118 | 1.05e-07 |  |  |
| female flowering | outlier | 1 | 9.94e-08 | 9.94e-08 | 0.0264 | 0.871 |
|  | frequency bin | 9 | 0.00197 | 0.000218 | 58.1 | 1.35e-106 |
|  | outlier:frequency bin | 9 | 0.000572 | 6.35e-05 | 16.9 | 3.48e-28 |
|  | Residuals | 113000 | 0.424 | 3.76e-06 |  |  |
| fusarium | outlier | 1 | 1.11e-07 | 1.11e-07 | 0.0798 | 0.778 |
|  | frequency bin | 9 | 0.000266 | 2.95e-05 | 21.1 | 4.13e-36 |
|  | outlier:frequency bin | 9 | 3.61e-05 | 4.01e-06 | 2.87 | 0.00217 |
|  | Residuals | 113000 | 0.157 | 1.4e-06 |  |  |
| grain yield | outlier | 1 | 5.07e-05 | 5.07e-05 | 1.48 | 0.224 |
|  | frequency bin | 9 | 0.0097 | 0.00108 | 31.5 | 1.1e-55 |
|  | outlier:frequency bin | 9 | 0.00344 | 0.000382 | 11.2 | 1.31e-17 |
|  | Residuals | 113000 | 3.86 | 3.42e-05 |  |  |
| oil content | outlier | 1 | 9.52e-06 | 9.52e-06 | 23.7 | 1.12e-06 |
|  | frequency bin | 9 | 2.29e-05 | 2.55e-06 | 6.35 | 4.75e-09 |
|  | outlier:frequency bin | 9 | 3.49e-05 | 3.87e-06 | 9.65 | 6.95e-15 |
|  | Residuals | 113000 | 0.0452 | 4.01e-07 |  |  |
| plant height | outlier | 1 | 0.000502 | 0.000502 | 10.5 | 0.00119 |
|  | frequency bin | 9 | 0.033 | 0.00367 | 76.8 | 1.44e-142 |
|  | outlier:frequency bin | 9 | 0.0124 | 0.00138 | 28.9 | 9.98e-51 |
|  | Residuals | 113000 | 5.38 | 4.78e-05 |  |  |
| protein content | outlier | 1 | 8.13e-07 | 8.13e-07 | 2.36 | 0.124 |
|  | frequency bin | 9 | 0.000184 | 2.04e-05 | 59.3 | 5.19e-109 |
|  | outlier:frequency bin | 9 | 3.04e-05 | 3.37e-06 | 9.8 | 3.76e-15 |
|  | Residuals | 113000 | 0.0388 | 3.44e-07 |  |  |

**Figure S2.**
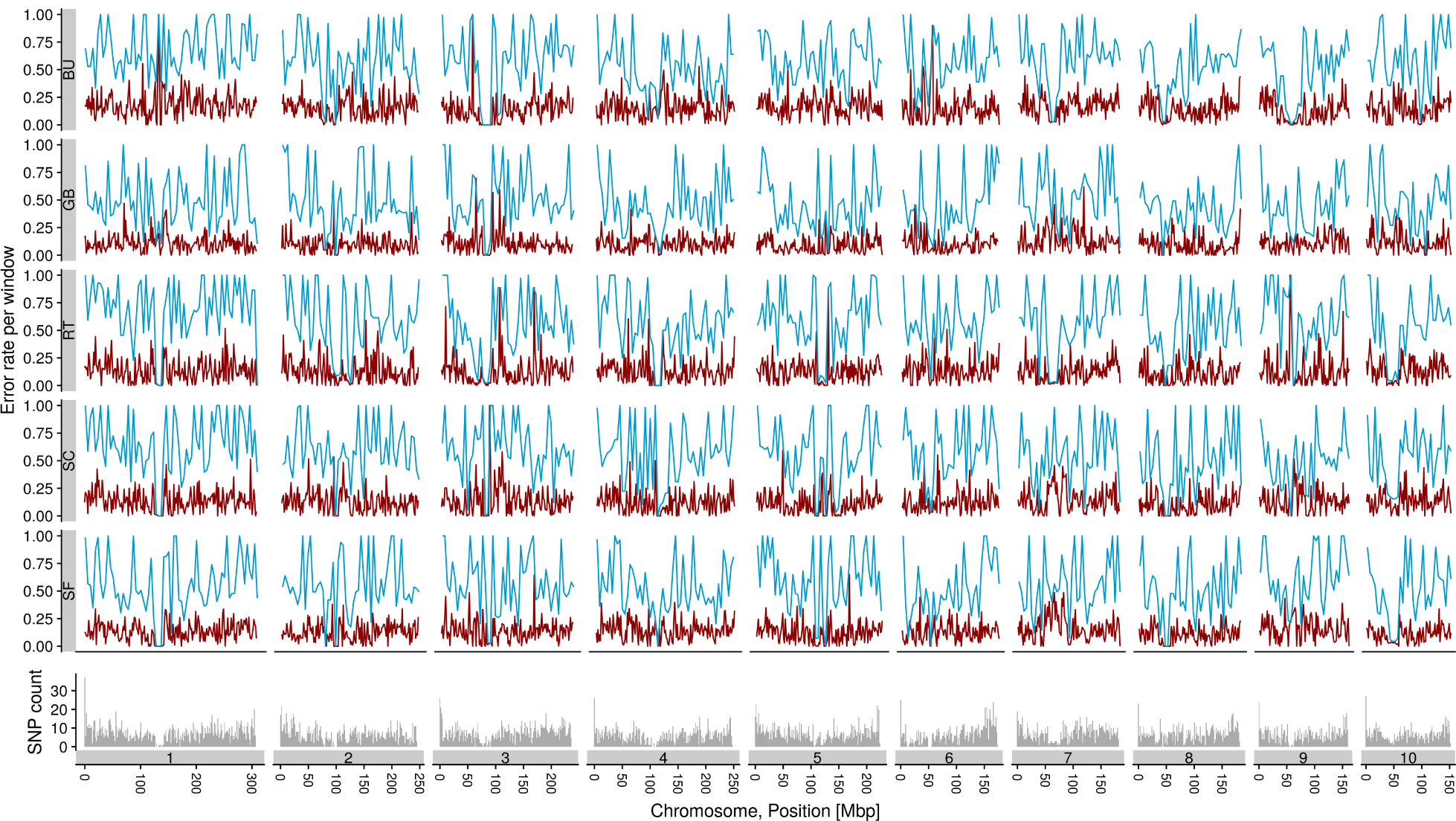
Imputation error rate for DH lines in five accessions. 10,000 known random SNPs were dropped and imputed to compute the error rate represented by mean error in 1.5 Mbp window (red line) and maximum error in 4.5 Mbp window (blue line). SNP density of the Illumina chip in 1.5 Mbp windows shown in bottom panel.

**Figure S3.**
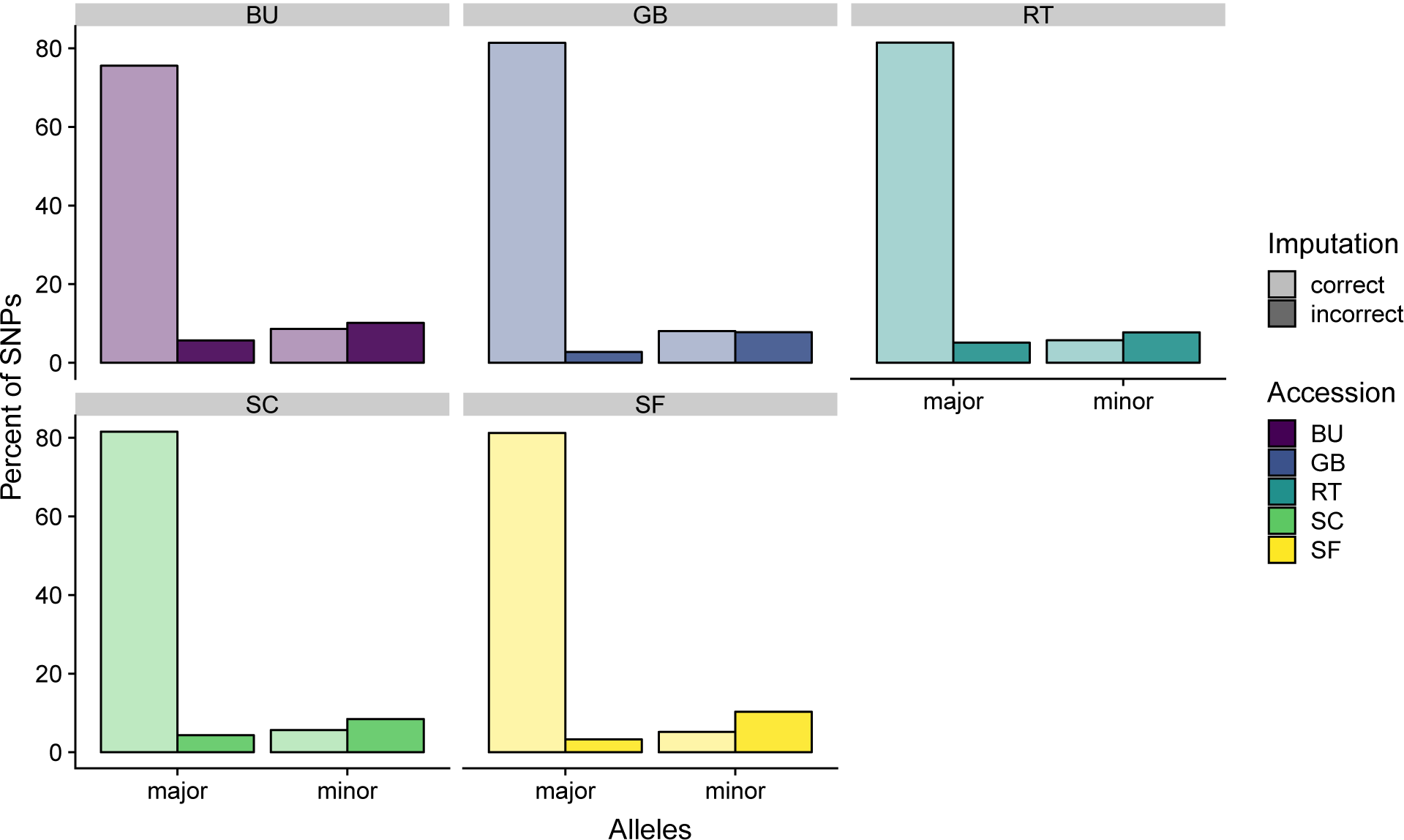
Percentages of correctly and incorrectly imputed SNPs in the imputation test runs.

**Figure S4.**
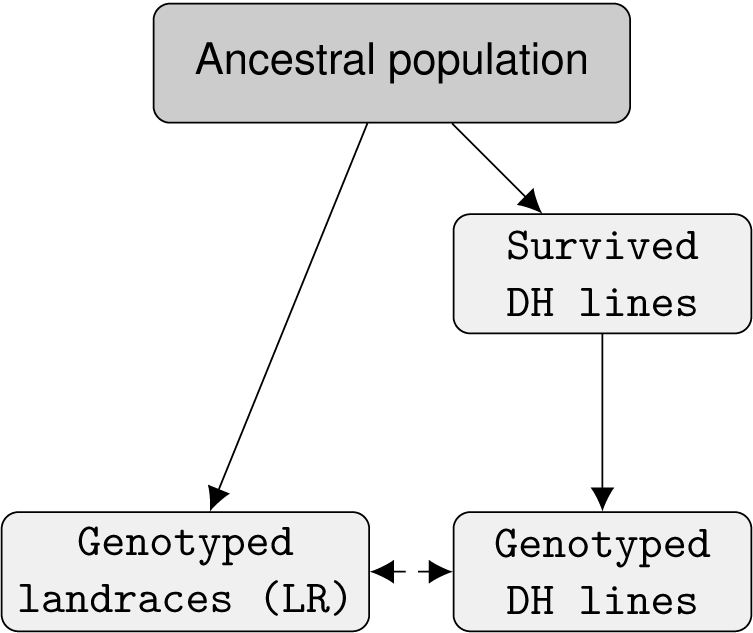
Assumed, simplified sampling structure for DH and LR we used to calculate ancestral frequencies and *p*-values.

**Figure S5.**
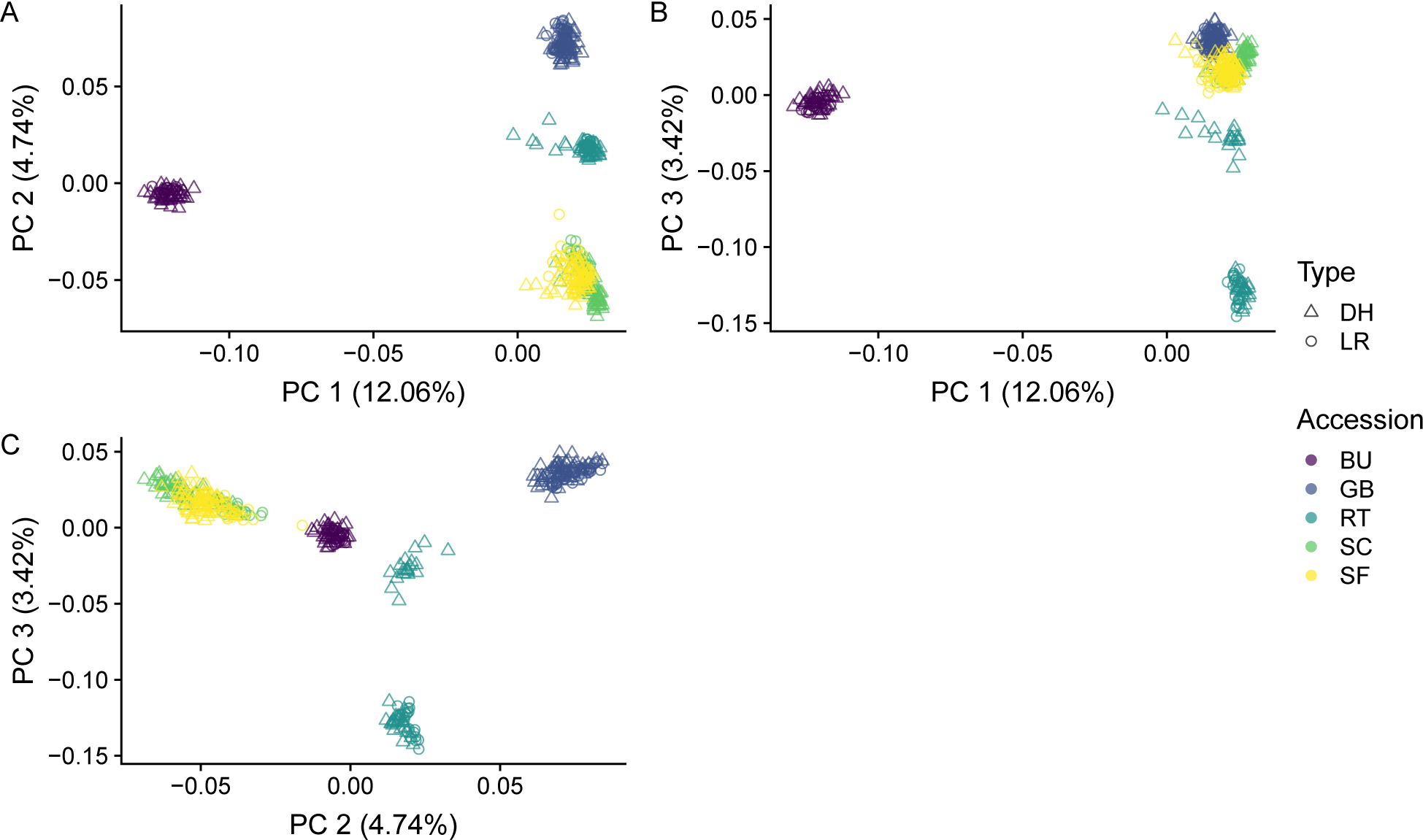
Principal component analysis for DH and LR of the 50k dataset, plot of principal component 1 and 2 (A) shows common clusters for LR and DH in respective accessions. However, principal component 3 separates the DH set of accession RT (B, C).

**Figure S6.**
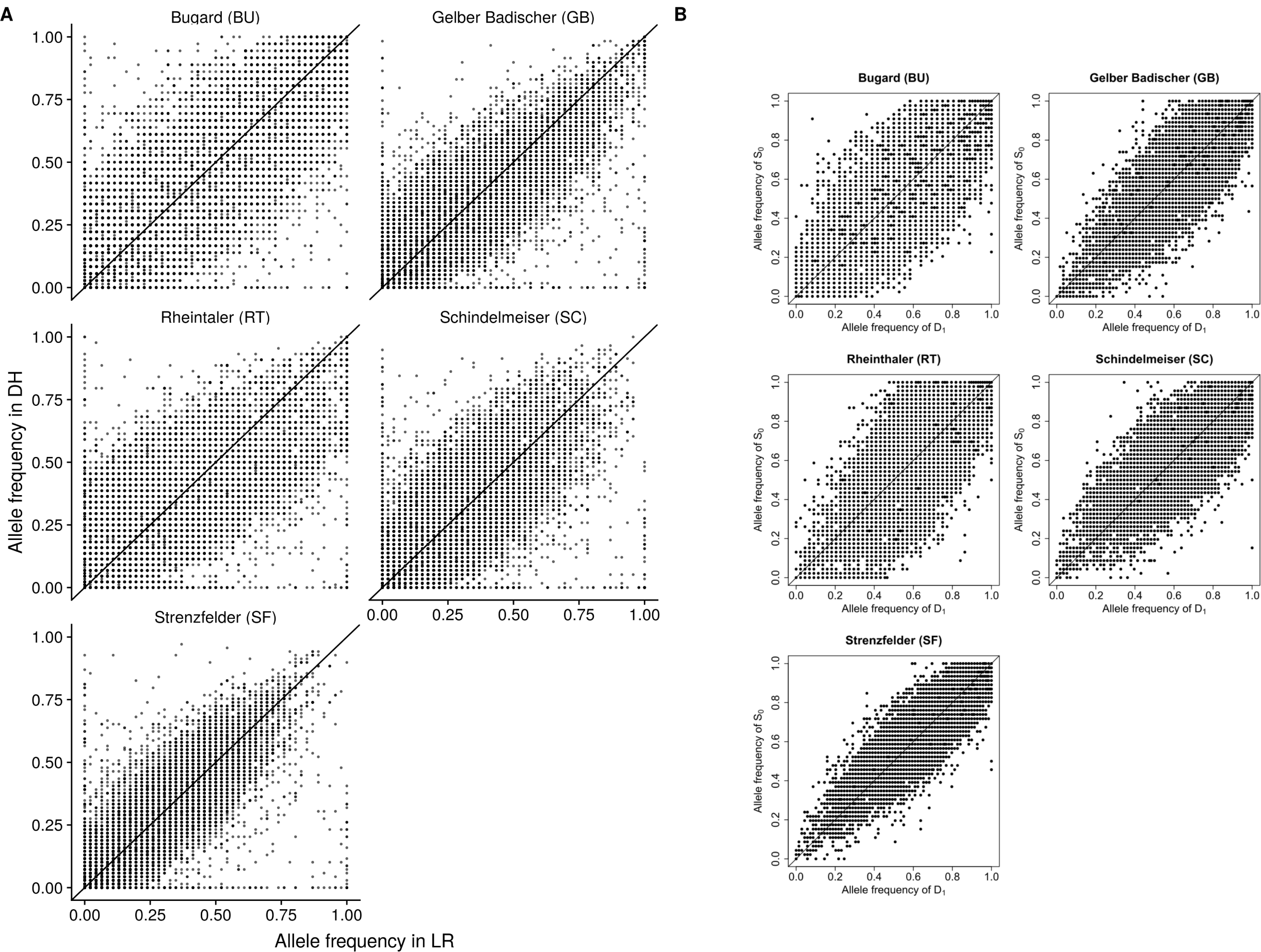
Comparison of joint frequency spectra with published data. (A) Landrace population and DH line alternative allele frequencies estimated from filtered supplementary dataset from Melchinger *et al*. (2017). (B) For comparison published figure (Melchinger *et al.* 2017).

**Figure S7.**
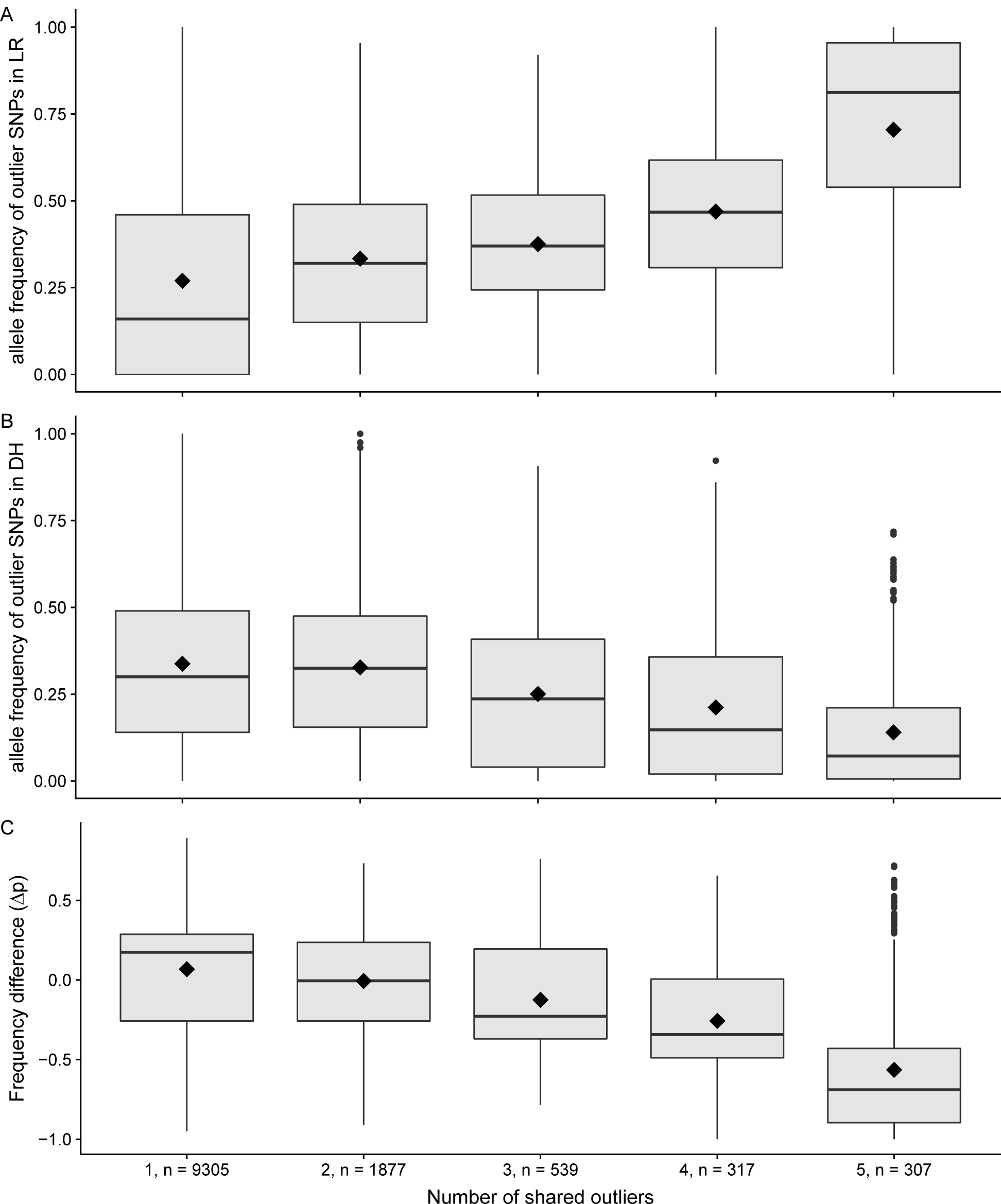
Mean allele frequencies in LR populations (A) and DH lines (B) calculated per number of shared outlier alleles. Outlier alleles, that are shared more often across populations are more likely to have low frequencies and to be lost, while unique outliers change only little in frequency (C). Numbers on x axis ticks correspond to number of shares and number of alleles in this column.

**Figure S8.**
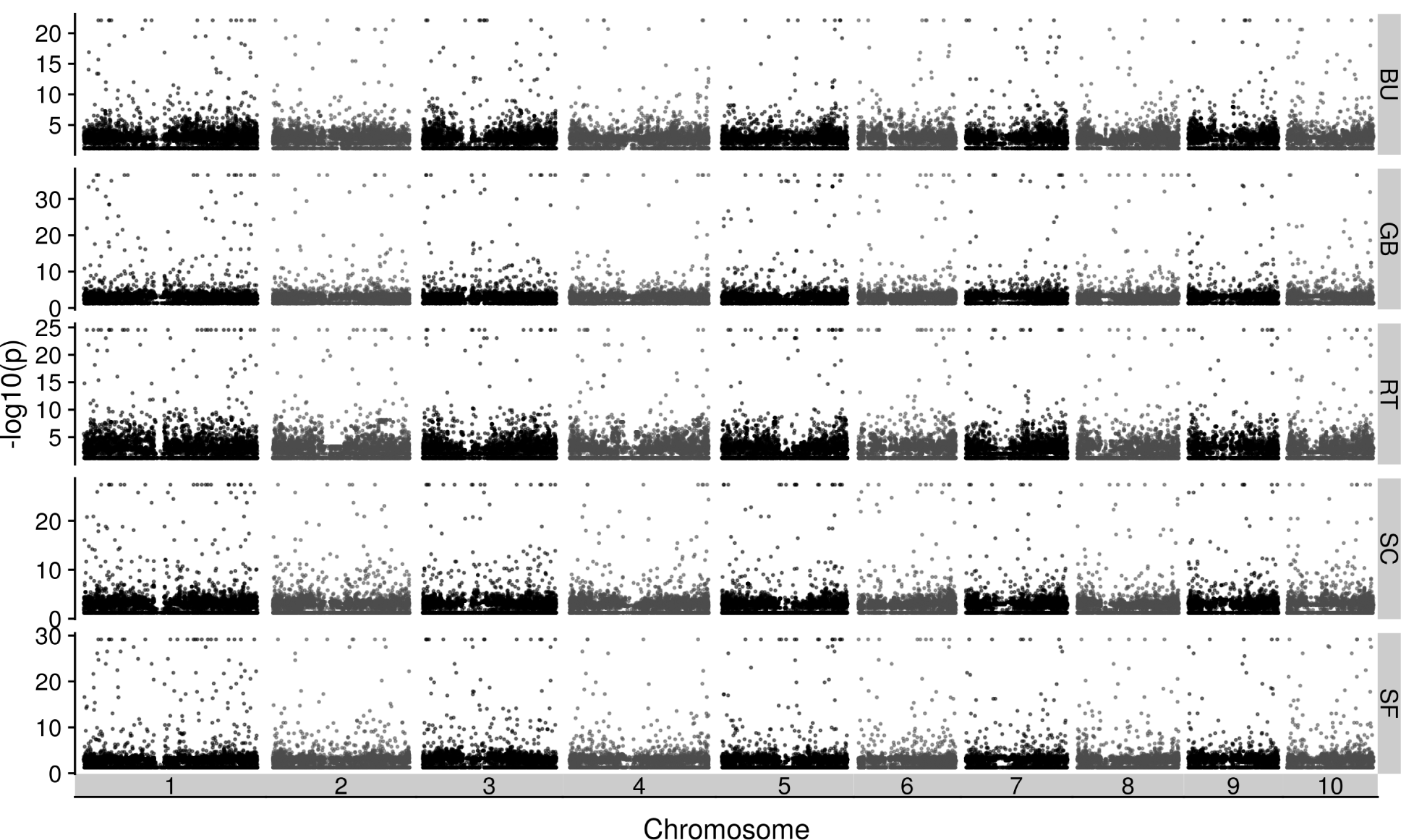
Joint probabilities of DH and LR allele frequency for all accessions and chromosomes.

**Figure S9.**
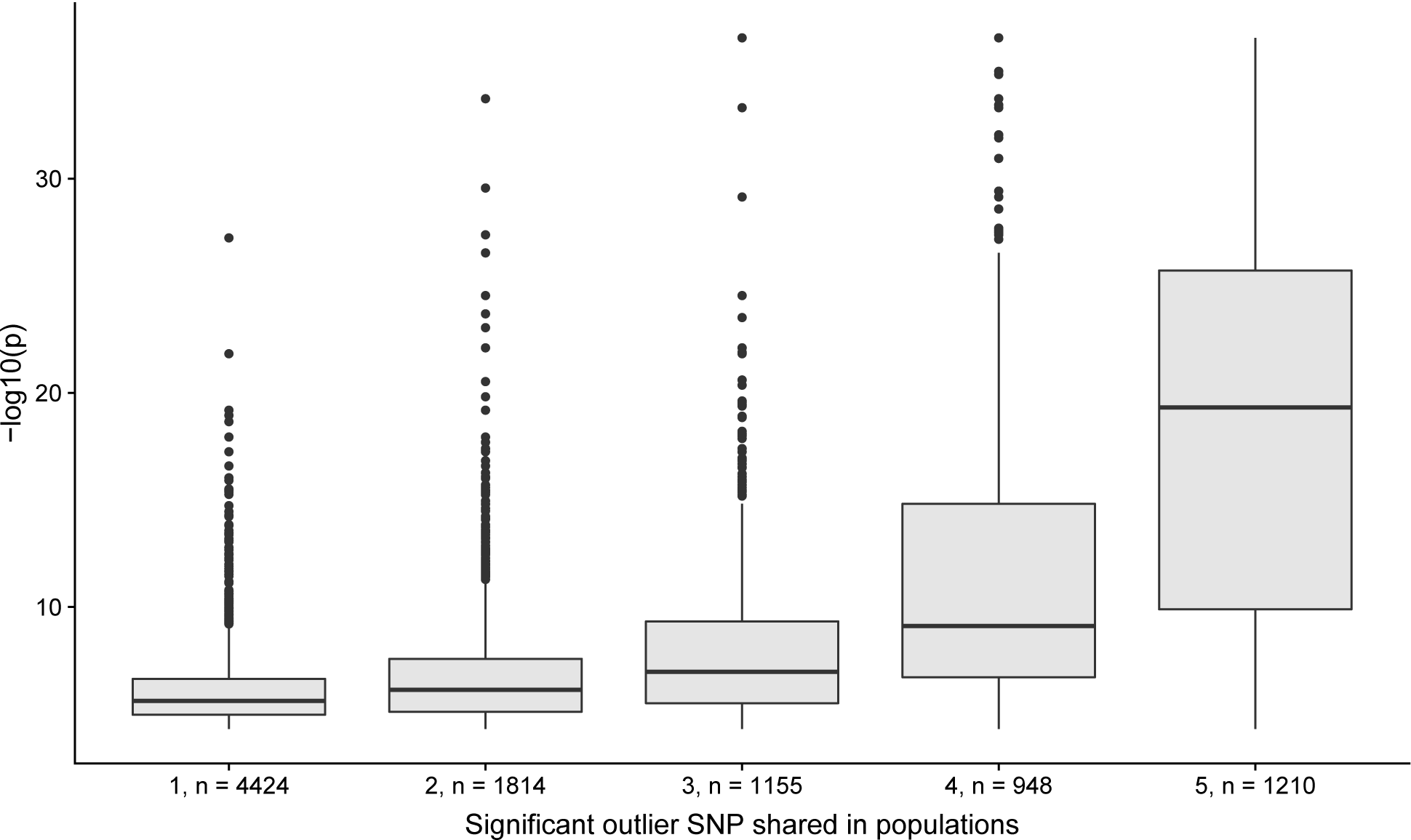
Shared significant –*log*_10_(*p*) values of the probability test and their overlaps among accessions show that high values are found primarily in frequently shared outlier SNPs. Numbers on x-axis correspond to the shared populations and number of outlier-SNPs in this class.

**Figure S10.**
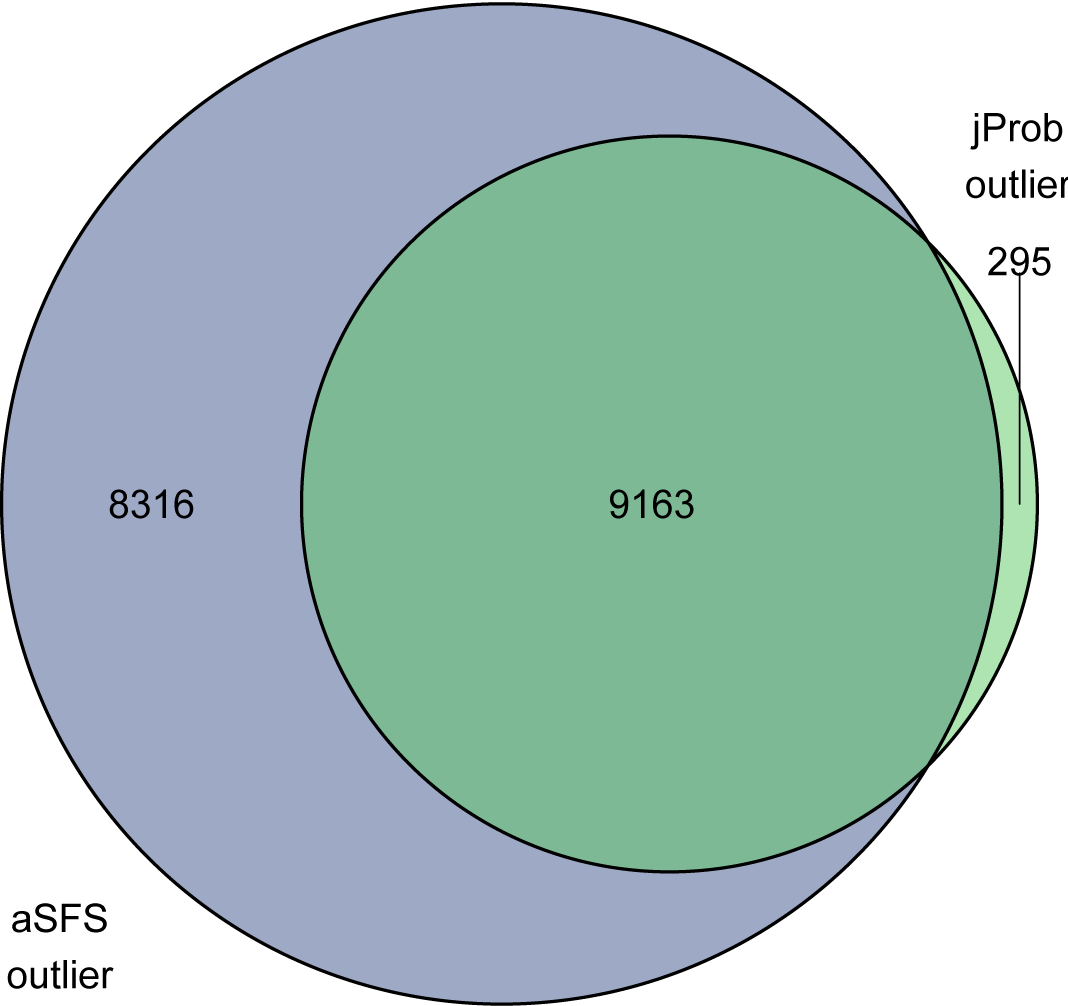
Shared outliers of aSFS and joint probability (jProb) tests, numbers in circles refer to summarized number of outlier in category.

**Figure S11.**
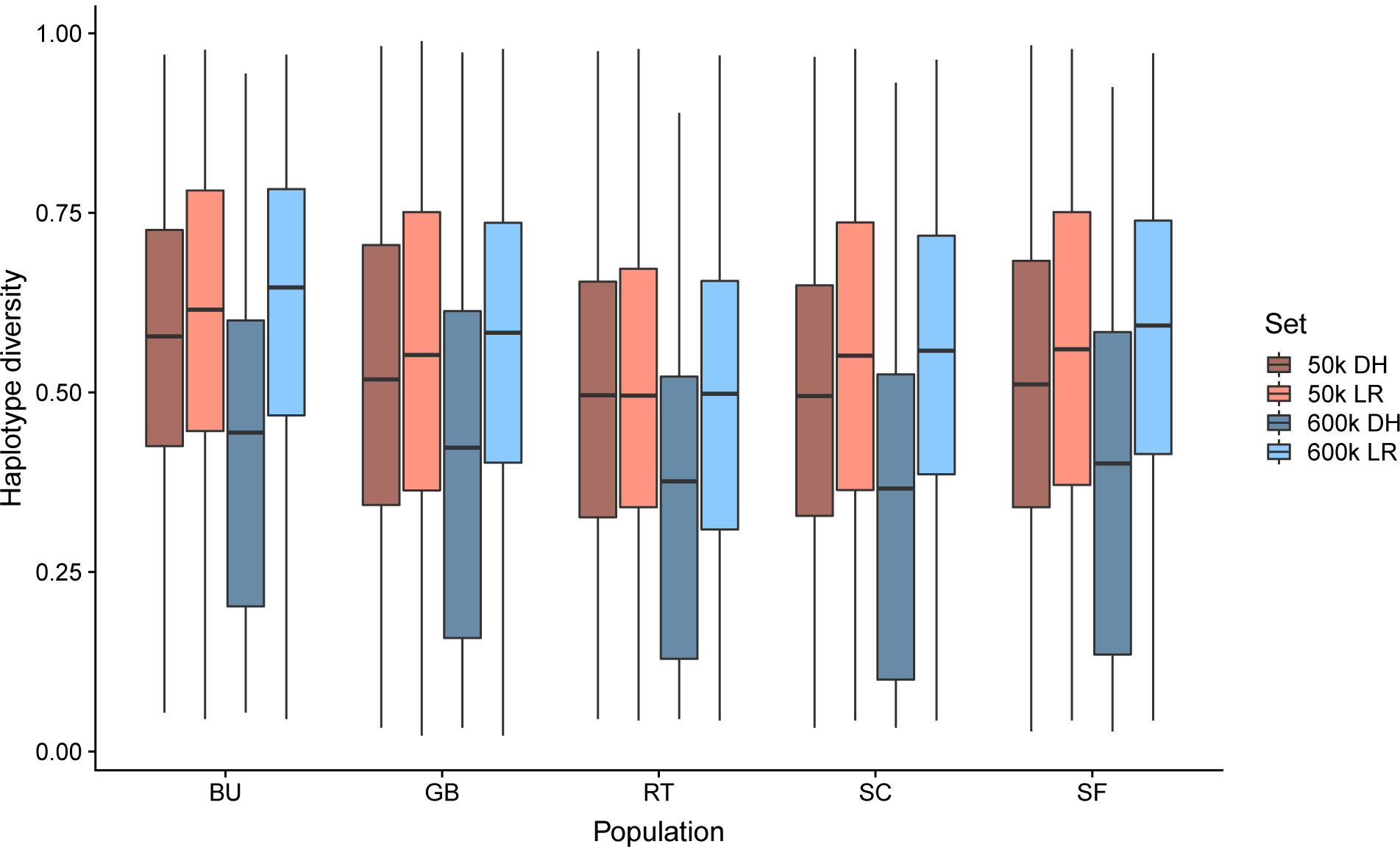
Comparison of haplotype diversity in 50k and 600k datasets in 50kb windows with more than one haplotype shows increase in haplotype diversity in the 600k LR dataset compared to the 600k DH, as well as a reduction of 50k LR compared to 600k DH. This difference is not visible in the DH, indicating that some diversity is missed during the imputation.

**Figure S12.**
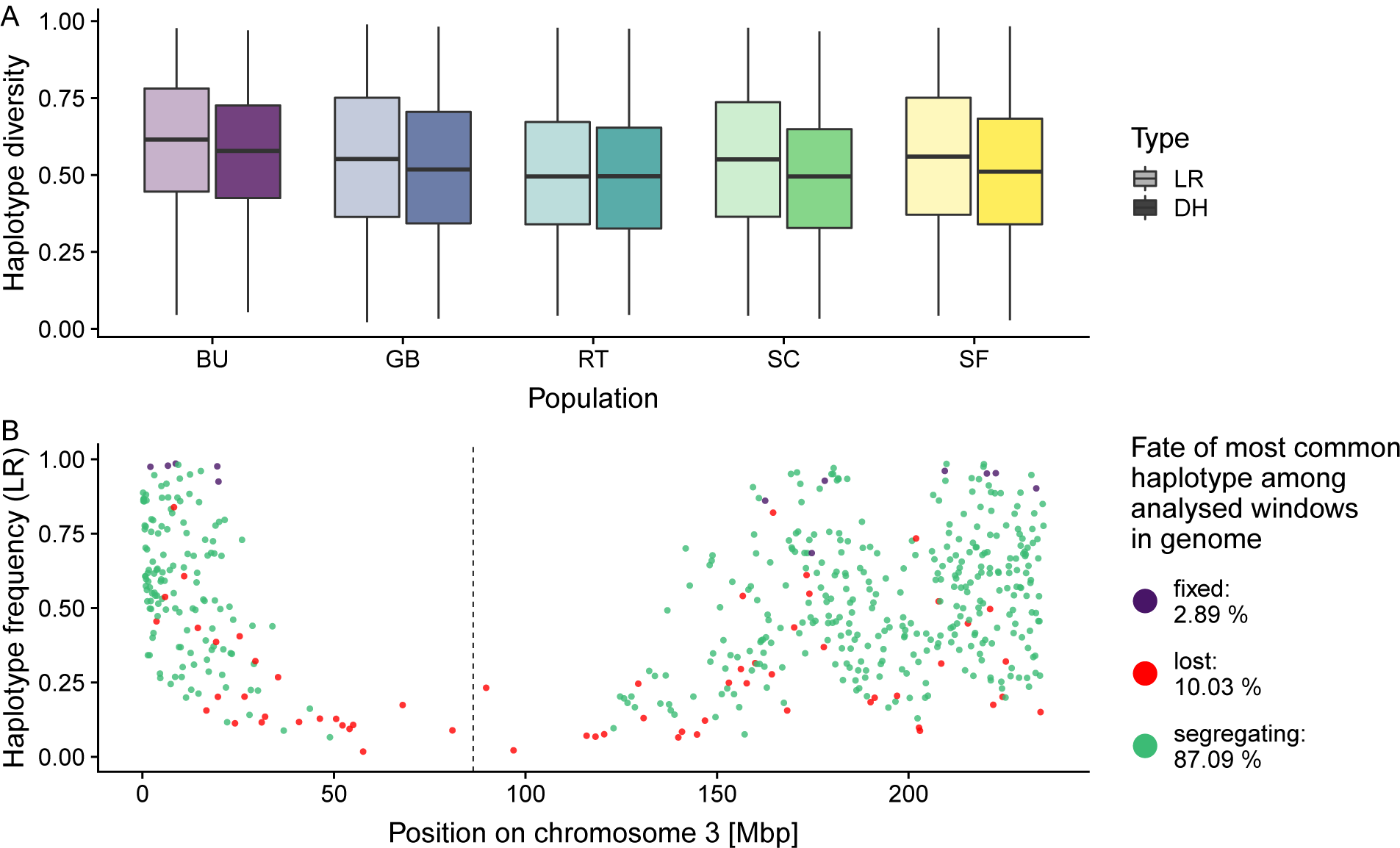
Similar to Figure 3 we analysed haplotypes of the 50k data using windows based on genetic distance (0.2 cM windows). (A) The reduction in haplotype diversity is less pronounced due to higher ascertainment bias and lower SNP density of the 50k chip and imputational error of the 600k DH. (B) Fixed major haplotypes occur very rarely, lost haplotype, however, quite frequently. Reduced SNP density and differences in ascertainment panel sizes between 50k and 600k arrays remove the signal of the putative inversion.

**Figure S13.**
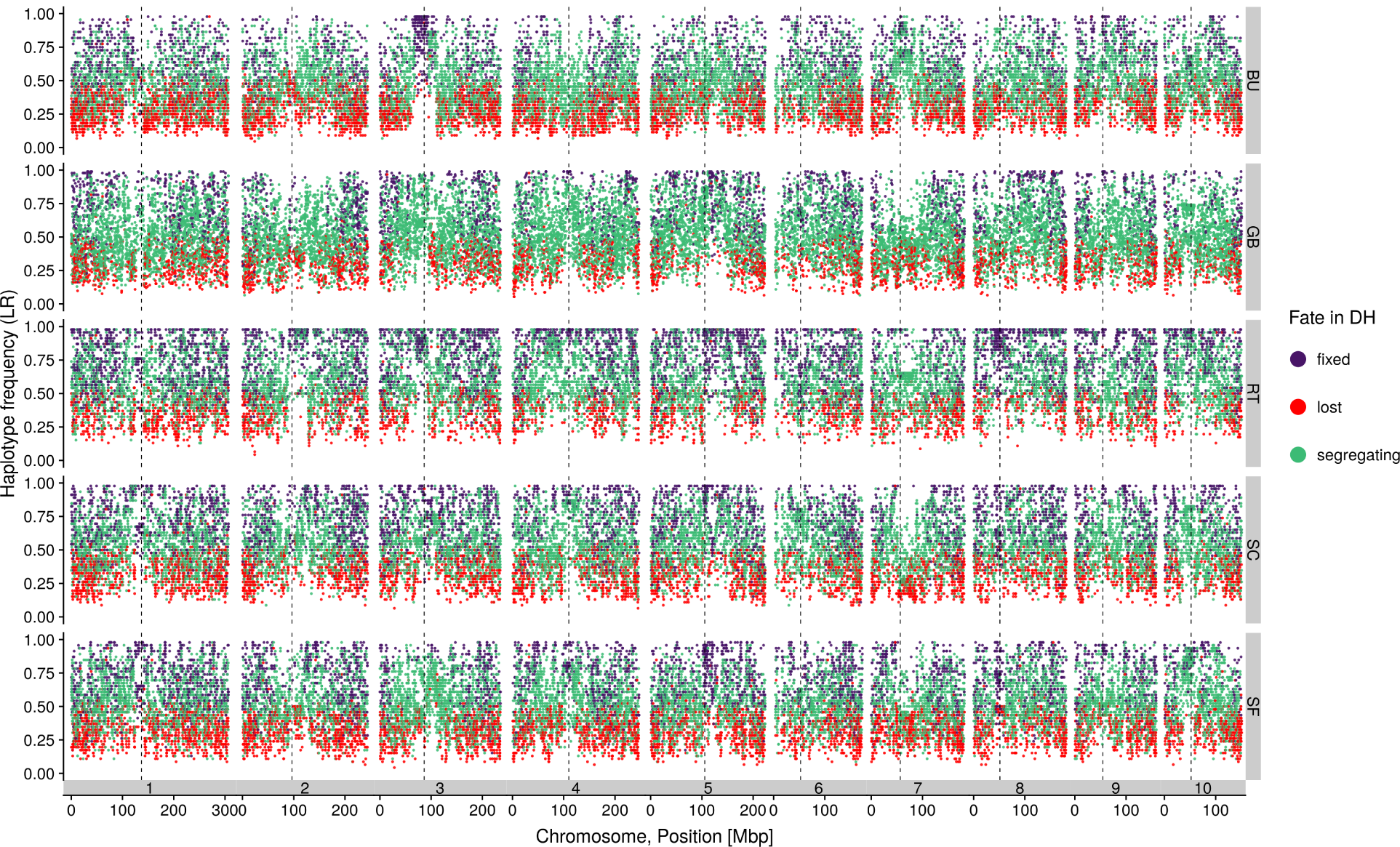
Fate of the most common haplotypes in all accessions. Centromeres are shown as dashed lines.

**Figure S14.**
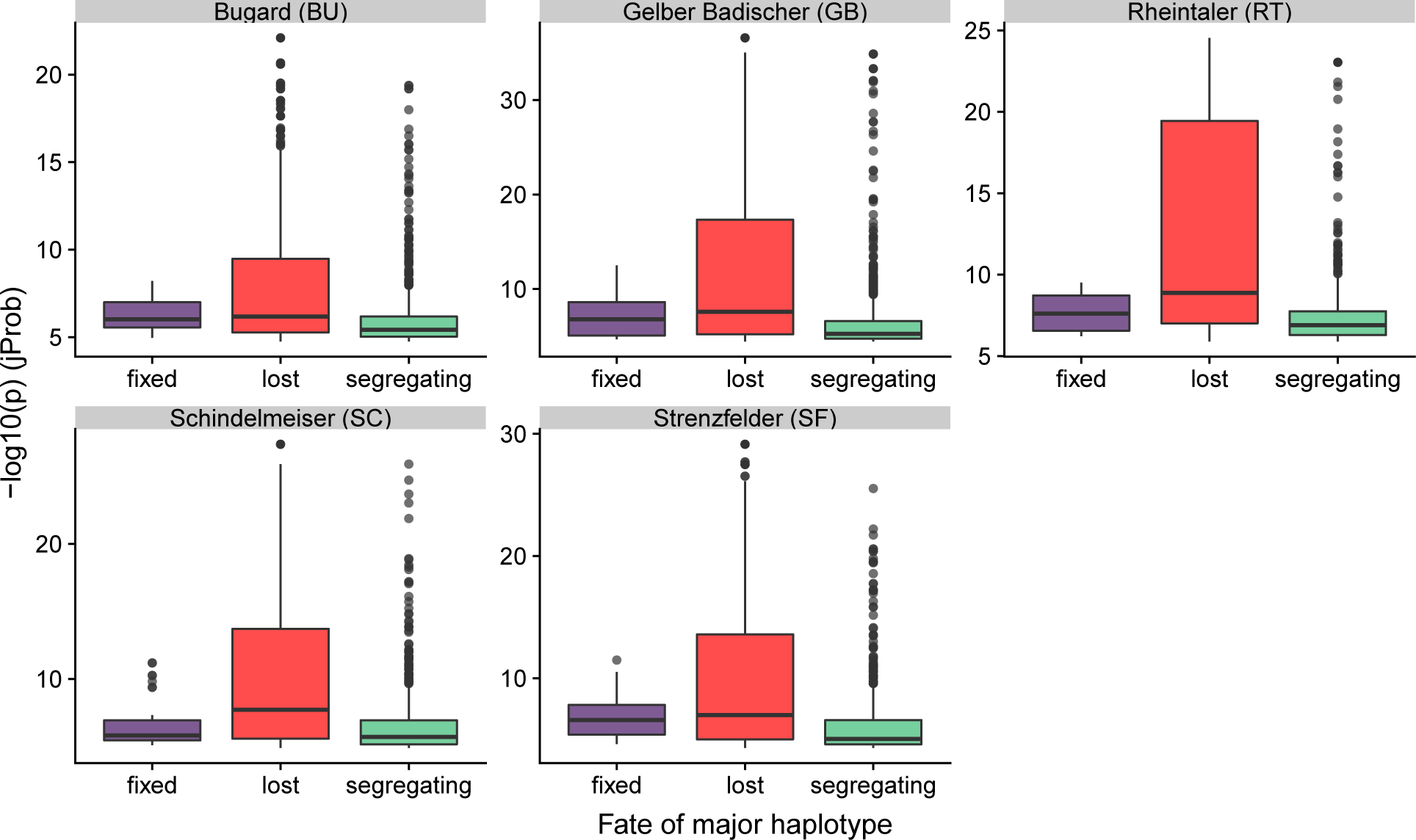
Fate of major haplotypes in 50kb windows outliers from the joint probability test reveal the highest significance levels for outlier in regions with large scale losses of haplotypes.

**Figure S15.**
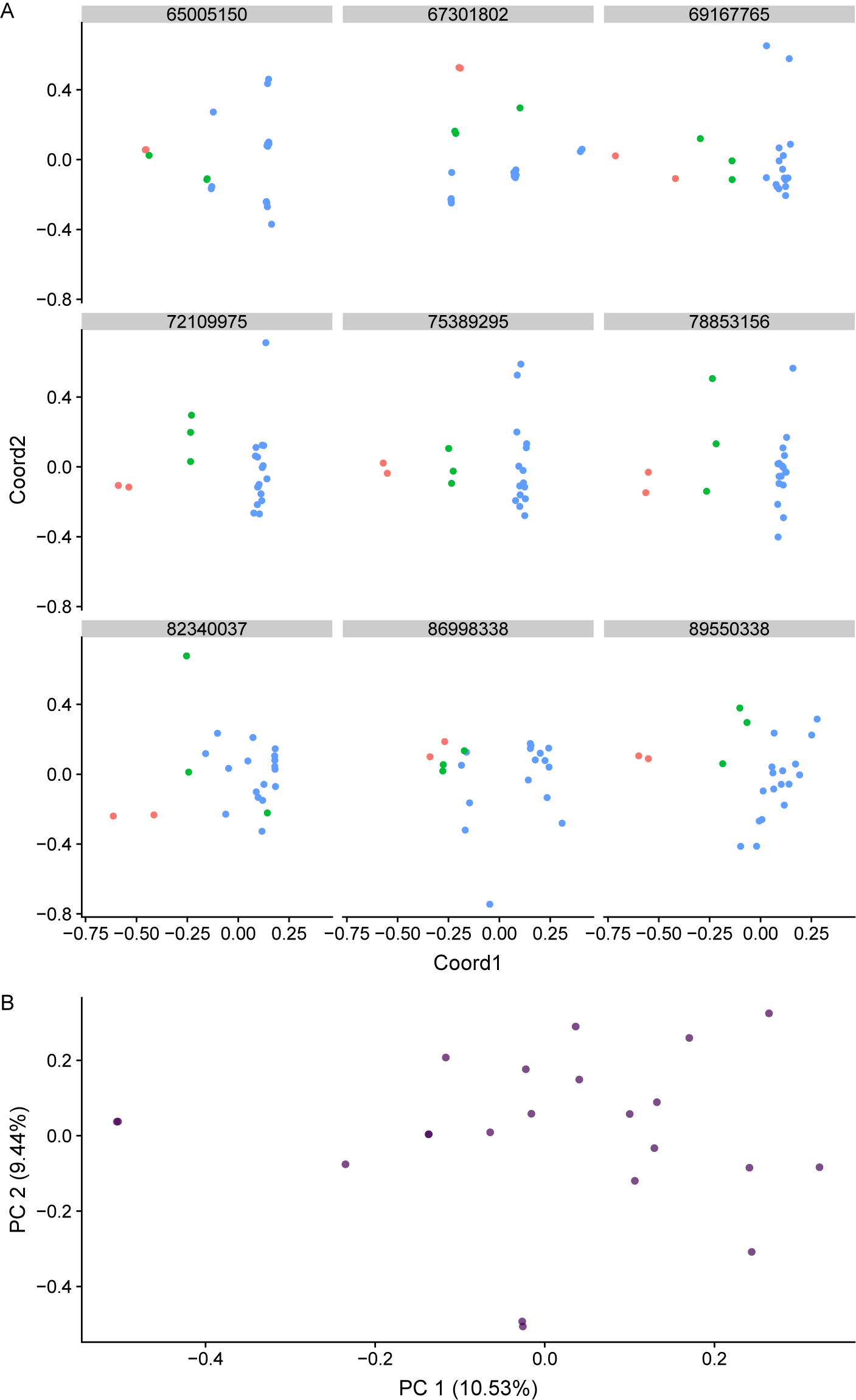
(A) Local PCA reveals structural variation in multiple consecutive windows (chromosome 3, 72,109,975; 75,389,295; 78,853,156) in putative inversion region of accession BU. Facet labels correspond to window start positions. Each windowed PCA was computed using 500 SNPs of the 600k LR BU dataset. Color codes refer to 3 clusters observed in window 72,109,975 to track clustering patterns in adjacent windows. (B) No structure is observed in principle components computed for genome-wide 600k data of accession BU (LR).

**Figure S16.**
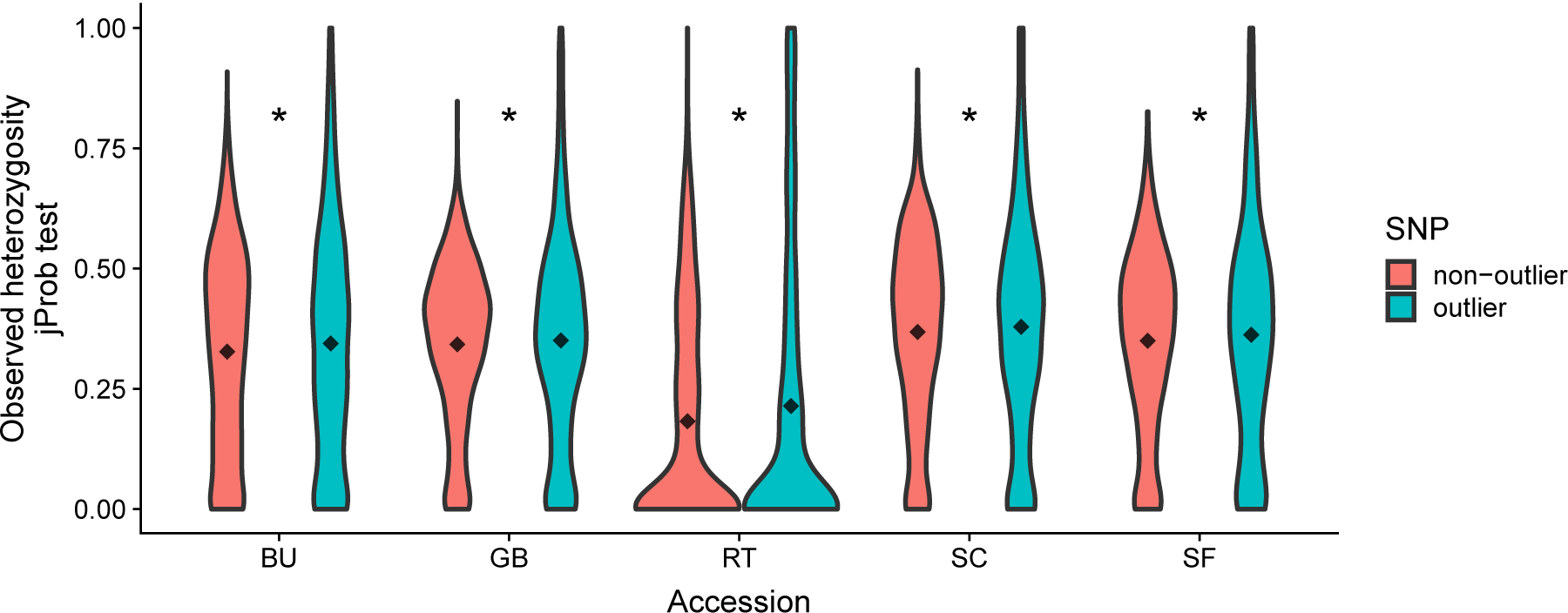
Violin plots for the frequencies of heterozygous genotypes of LD-pruned non-outlier SNPs and outlier SNPs in LR accessions for joint probability outlier. Diamonds indicate group means. Comparisons with asterisks have significantly different means (*p <* 0.05).

**Figure S17.**
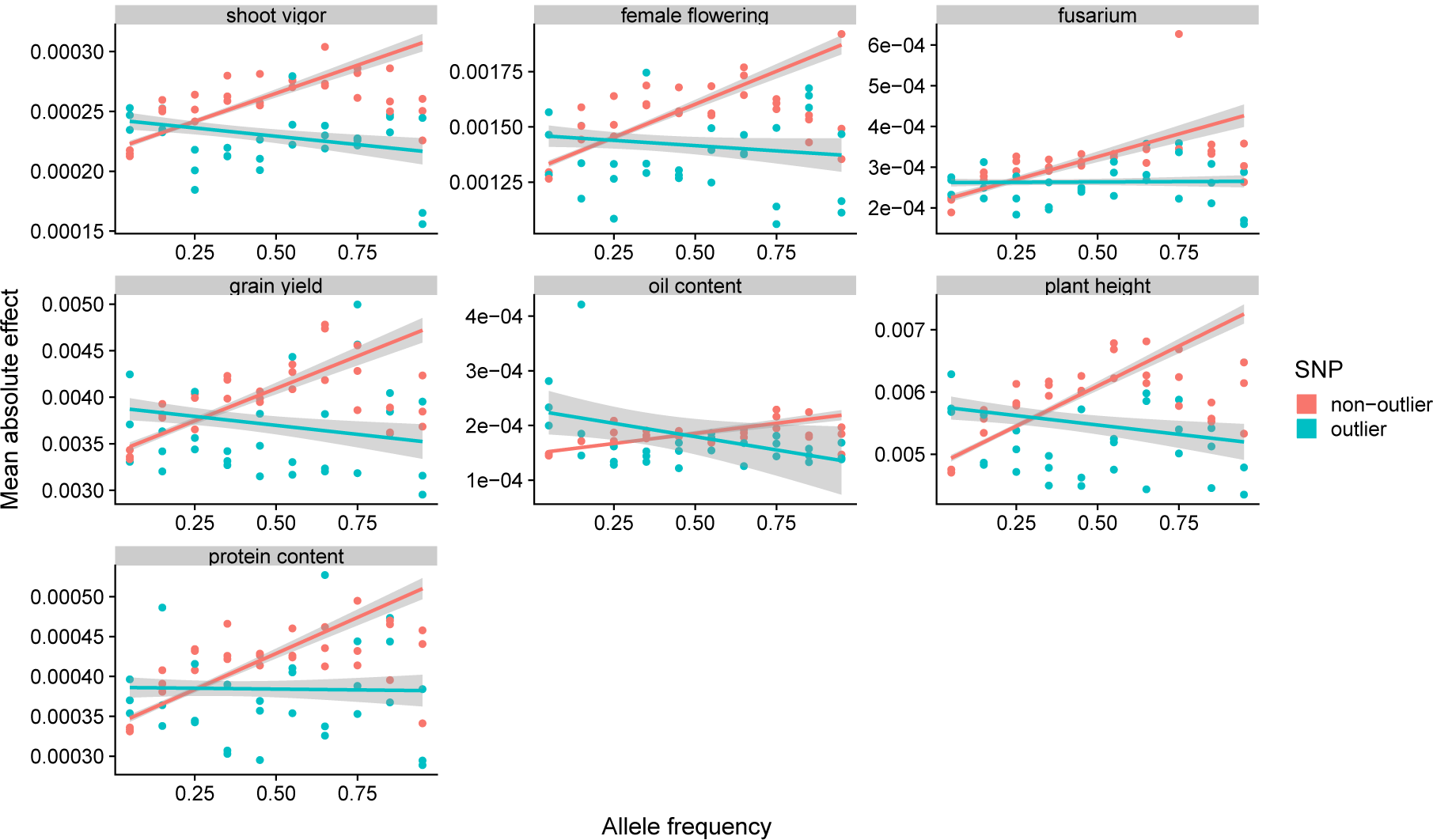
Comparison of mean absolute effect sizes (*α*) for outliers and non-outliers in different frequency bins in three populations used in GWAS. By fitting a linear regression model 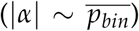 for fitness-related traits and outlier status we explore significant interactions in ANOVA (Table S6) between outlier status and frequency bin. Regression slopes with different signs highlight the deleterious action of outlier SNPs. Shaded areas around regression lines are 95% confidence intervals.

**Figure S18.**
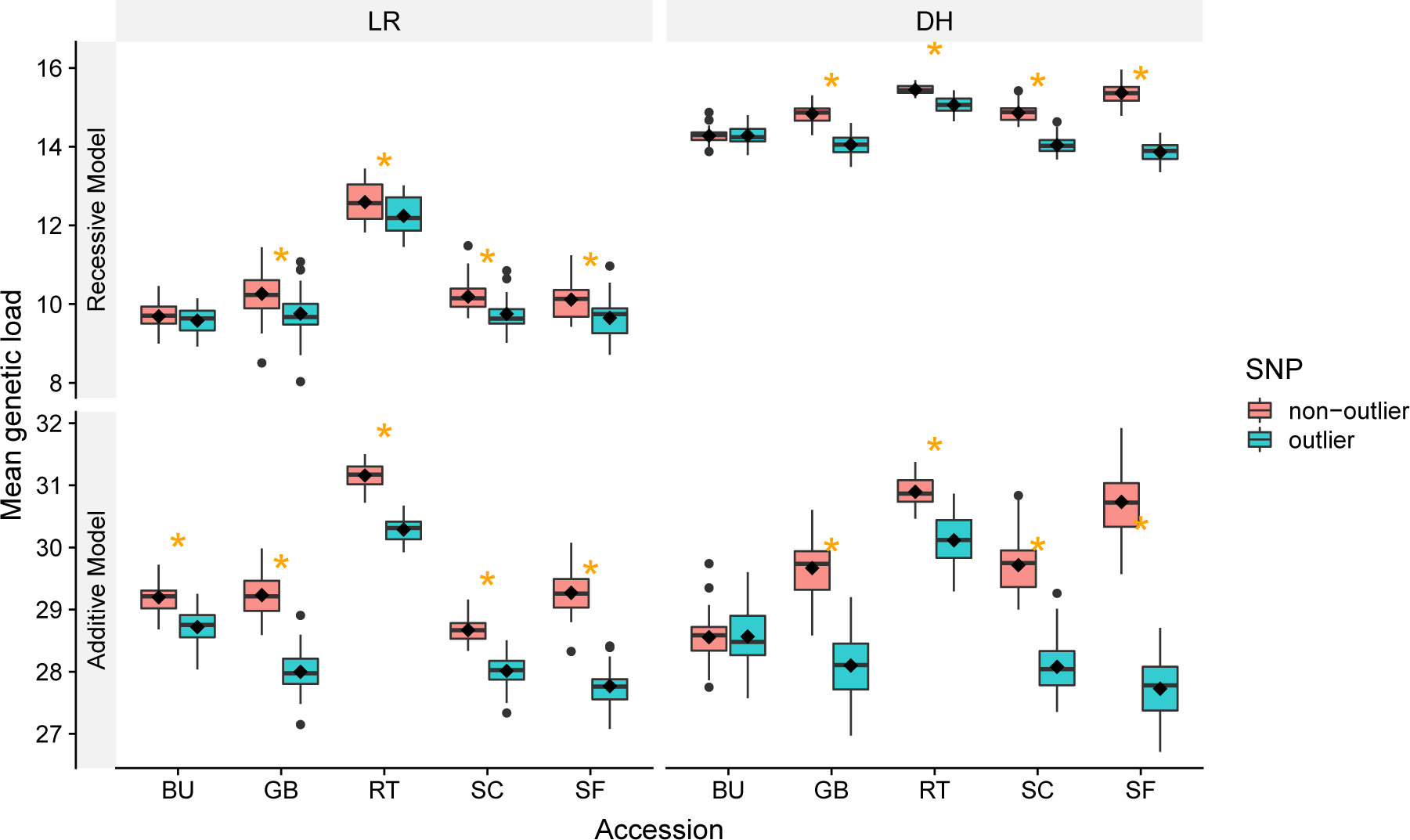
Mean individuals’ GERP sum in 1 cM region for SNPs with GERP > 0 per DH-LR pair and SNP-type reveal differences in putative genetic load comprised within accessions and populations for the additive and recessive model. Group means are represented by diamonds. Orange asterisks mean significantly different outlier/non-outlier means within accession and population (t-test, *p <* 0.05).

